# Feature-specific divisive normalization improves natural image encoding for depth perception

**DOI:** 10.1101/2024.09.05.611536

**Authors:** Long Ni, Johannes Burge

## Abstract

Vision science and visual neuroscience seek to understand how stimulus and sensor properties limit the precision with which behaviorally-relevant latent variables are encoded and decoded. In the primate visual system, binocular disparity—the canonical cue for stereo-depth perception—is initially encoded by a set of binocular receptive fields with a range of spatial frequency preferences. Here, with a stereo-image database having ground-truth disparity information at each pixel, we examine how response normalization and receptive field properties determine the fidelity with which binocular disparity is encoded in natural scenes. We quantify encoding fidelity by computing the Fisher information carried by the normalized receptive field responses. Several findings emerge from an analysis of the response statistics. First, broadband (or feature-unspecific) normalization yields Laplace-distributed receptive field responses, and narrowband (or feature-specific) normalization yields Gaussian-distributed receptive field responses. Second, the Fisher information in narrowband-normalized responses is larger than in broadband-normalized responses by a scale factor that grows with population size. Third, the most useful spatial frequency decreases with stimulus size and the range of spatial frequencies that is useful for encoding a given disparity decreases with disparity magnitude, consistent with neurophysiological findings. Fourth, the predicted patterns of psychophysical performance, and absolute detection threshold, match human performance with natural and artificial stimuli. The current computational efforts establish a new functional role for response normalization, and bring us closer to understanding the principles that should govern the design of neural systems that support perception in natural scenes.

## Introduction

Understanding how perceptual systems respond to natural signals is a topic of enduring interest in vision and neuroscience research. Although the topic has been recognized as having fundamental importance since the advent of modern vision science, only in the past twenty years have the experimental and computational tools matured sufficiently so that substantial progress can be made (Simoncelli & Olshausen 2001; Geisler, 2008; Geisler & Ringach, 2009; Burge, 2020). Progress has been achieved using multiple methods, ranging from direct neurophysiological measurement to computational modeling. Characterizing how receptive fields are driven by natural images is important to this effort. Knowledge of how responses are altered by different receptive field properties, and other aspects of sensory processing, is critical for understanding designs of neural systems that maximize performance (Geisler, 1989; Olshausen & Field, 1996; Burge, 2020). Quantifying the statistical properties of receptive field response to natural signals can help place data-constrained, image-computable models of visual information processing on firm, ecologically-valid foundations.

A primary function of our visual system is to estimate and categorize behaviorally relevant latent variables from images of the natural environment. To do so, image features that carry information about those latent variables must be extracted from the retinal images. Determining the image features that receptive fields should select for, and how the extracted features should be combined, are difficult problems.

The premise of this article is that an examination of the statistical properties of receptive field responses to natural stimuli that signal different values of behaviorally-relevant latent variables will help one gain a deeper understanding of principles underlying the computations that neural systems perform, and of the limits of perceptual performance. The computations that support the optimal computations—e.g., optimal combination of the receptive field responses—are dictated by the statistical relationship between the features selected for by useful receptive fields and the latent variable(s) of interest. Measuring how natural stimulus variability—an ecologically valid form of ‘nuisance’ variability—and how aspects of the response model shape the statistics of receptive field response is an obvious starting place.

Here, using stereo-depth perception as a model system, we report an extensive computational analysis of how binocular receptive fields are driven by natural stereo-images. The analyses are focused on understanding i) how properties of the response model impact the response statistics, and ii) how those statistics should constrain and shape performance in stereo-(i.e. binocular-disparity-based-) depth perception tasks.

The article is organized as follows. First, we examine how binocular receptive fields respond to natural stereo-images. We quantify how different forms of response normalization—broadband (feature-unspecific) and narrowband (feature-specific) normalization—alter the amount of Fisher information about disparity in the responses. Second, we examine how responses are modulated by fixation disparity, and how well fixation disparity can be discriminated from the responses of individual receptive fields with different spatial frequency preferences. Finally, we analyze how pairs and populations of responses are modulated by fixation disparity, and use these response statistics to predict patterns of human disparity-discrimination performance with natural stereo-images.

To preview the major results, we find that the form of normalization and the spatial frequency selectivity of the receptive fields strongly impacts the amount of Fisher information about the latent variable in their responses: coding fidelity is higher with narrowband (feature-specific) than with broadband (feature-unspecific) normalization. We find that this functional advantage increases systematically as receptive field population size increases. And we find that distinctive patterns of psychophysical performance are predicted by the statistics of response to natural images.

## Results

The analyses described in this section are focused on understanding how canonically-shaped binocular receptive fields respond to natural stereo-images, and how the response properties shape and constrain performance in stereo-depth perception. First, we describe the natural stereo-image dataset and the binocular receptive field response model on which the analyses are based. Next, we present response statistics, encoding fidelity, and discrimination performance based on responses of individual receptive fields with different spatial frequency preferences, pairs of receptive fields, and an entire receptive field population. Throughout, specific emphasis is placed on how the form of response normalization changes coding fidelity with receptive field population size. Although the results described here are specific to stereo-depth perception, there is evidence that the principles described here should be general to a range of other sensory-perceptual tasks—for example, focus error (Burge & Geisler, 2011; 2012), 2D motion (Burge & Geisler, 2015; Chin & Burge, 2020), and 3D motion estimation (Herrera & Burge, 2024).

### Natural-stereo images

Stereo-image patches of natural scenes were sampled from a dataset of stereo-images with co-registered laser-based distance measurements at each pixel (Burge, McCann, and Geisler 2016). Corresponding left- and right-eye-image points, and ground-truth relative disparities, were computed directly from the distance data assuming a virtual human observer with a 65mm interpupillary distance (Iyer & Burge, 2018; Fig. 1a).

**Figure 1:**
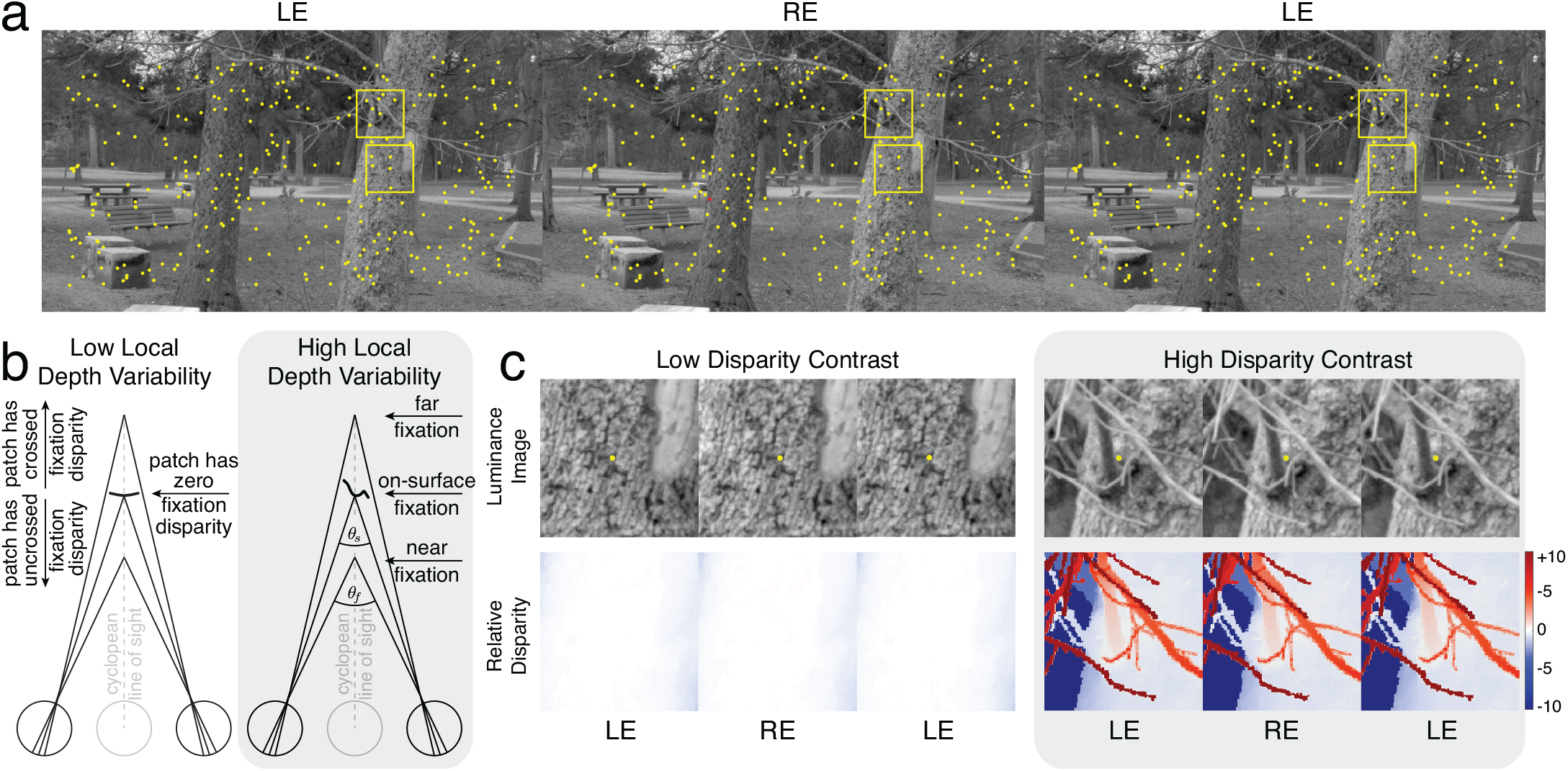
Fixation disparity and nuisance variability in natural stereo-images. **(a)** Natural stereo-image. Divergently-fuse the left two images, or cross-fuse the right two images to see the scene in stereo-3D. Corresponding points, which are computed from co-registered laser-based distance measurements at each pixel, are overlayed (yellow dots). Yellow boxes indicate example stereo-image patches (4/3ºx4/3º patches were analyzed in the paper; 2.5×2.5º patches are shown here for purposes of visualization). **(b)** For each local surface in the scene, left- and right-eye stereo-image patches were sampled with a fixation disparity (*δ*_0_ = *θ*_*t*_ − *θ*_*f*_)—roughly, how far a target surface point is from fixation (Howard & Rogers, 2002). Stereo-image patches were sampled from all scene locations. Some patches depicted scene regions with low local depth variability (left); others depicted regions with much more local depth variability (right). This article focuses its analysis on images depicting scene regions where local depth variability is low (< 2 arcmin), rather than high (grayed out). **(c)** Example stereo-image patches from the yellow boxes in (a). One patch has low local depth variability (and low disparity contrast; left). The other has high local depth variability (and high disparity contrast; right). The left- and right-eye intensity images (upper panels) and relative disparity maps computed from the distance measurements (lower panels) are both shown. The color bar shows pixel-wise relative disparities in arcmin. Disparity contrast is the root-mean-squared relative disparity with respect to a fixated corresponding point at the patch center (yellow dots in upper panel). This article focuses its analysis on stereo-images of flat (i.e. low disparity-contrast) surfaces.

Sampled stereo-image patches were centered on surface points in the scene (corresponding image points), offset by a fixation disparity that ranged from -30 to 30 arcmin (Fig. 1b). The local differences between the left- and right-eye patches defining each stereo-pair are due to two distinct factors: i) the imposed fixation disparity, and ii) the local depth structure of the surface(s) in the depicted scenes. Fixation disparity determines the overall leftward or rightward shift of the two images relative to one another. Local depth structure—which we quantify with disparity contrast (see *Methods*)— determines the local differences in the two images. For a flat frontoparallel surface straight ahead, for example, the left- and right-eye images are identical to one another except for a spatial shift. For a surface with a great deal of local depth variation, the left- and right-eye images will have many local differences between them (Fig. 1c). In this article, we analyze only approximately flat surfaces (i.e. surfaces with near-zero disparity contrast; see *Methods*). We do so because flat surfaces support the most precise disparity-based depth discriminations, and because research in the psychophysics and neuroscience literatures has typically been performed using flat surfaces. Future work will examine response statistics and estimation and discrimination performance associated with surfaces that are not flat.

Stereo-depth estimation is a difficult problem in part because of the many factors that inject random variation into binocular stereo-images. Luminance contrast-pattern variation--which is caused by differences in surface materials, textures, and lighting among other factors--is one source of nuisance variability that the nervous system must contend with. Local depth variation--which is due to the 3D structure of the surfaces composing scenes--is another (see Fig. 1bc). The distinct impact of these two sources of variability will be examined in future work (see *Discussion*).

Biological image systems must estimate the amount of fixation disparity at each point in a scene by determining how well local patches of the left- and right-eye images match one another. The consensus view is that this correspondence problem is solved with computations akin to a cross-correlation (Tyler & Julesz, 1978). In the nervous system, these computations depend on appropriate processing of receptive field responses to the left- and right-eye images (Fleet, Wagner, Heeger, 1996; Qian & Zhu, 1997; Anzai, Ohzawa, Freeman, 1999; Banks, Gepshtein, Landy, 2004; Burge & Geisler, 2014).

### Response Model

The impact of internal noise on perceptual performance has been studied intensively for decades. The manner in which natural nuisance variability impacts performance is less well understood (but see Hecht, Shlaer, Pirenne, 1942; Geisler, 1989; Burge & Geisler, 2015; Sebastian et al., 2017; Burge, 2020). Here, we study its impact using the classic computational framework of the linear-nonlinear subunit model.

Binocular receptive fields have left- and right-eye components. Each component is commonly modeled as having a Gabor-shaped weighting profile—a sinewave multiplied by a Gaussian—with a phase difference between the carrier wave of each component. Such receptive fields are said to select for phase disparity. It is also common to model the components as having the same Gabor-shaped weighting profile in offset positions from one another. Such receptive fields are said to select for position disparity. Both types occur in cortex (Cumming & DeAngelis, 2001), but evidence suggests that phase-disparity-selective receptive fields code for a broader range of disparities in primate and in cat (Prince, Cumming, Parker, 2002; Anzai, Ohzawa, Freeman, 1999). Although our analysis focuses on phase-disparity-selective receptive fields, all results are robust when position-disparity-selective receptive fields are included in the analysis, provided that the range of disparities coded by position-disparity-selective receptive fields is bounded, as it is in cortex.

We examine the response properties of receptive fields with spatial frequency preferences that range from 1-8 cycles per degree (Fig. 2ab; see *Methods*). The octave bandwidth of the receptive field component in each eye was 1.2 octaves and the orientation bandwidth was 42º, values that approximately match the median values recorded from simple cells in early visual cortex (De Valois, Albrecht, Thorell, 1982; Ringach, 2002). (All qualitative results are robust to biologically-realistic variation around these numbers, see Iyer & Burge, 2019.) At each spatial frequency, we consider the responses of a pair of receptive fields in quadrature (Adelson & Bergen, 1985). Quadrature receptive fields are orthogonal to one another 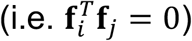 and select for the same region of the spatial frequency spectrum (i.e. have the same amplitude spectra). Receptive field pairs in quadrature phase extract all information in the region of the spatial frequency spectrum that the receptive fields select for.

**Figure 2.**
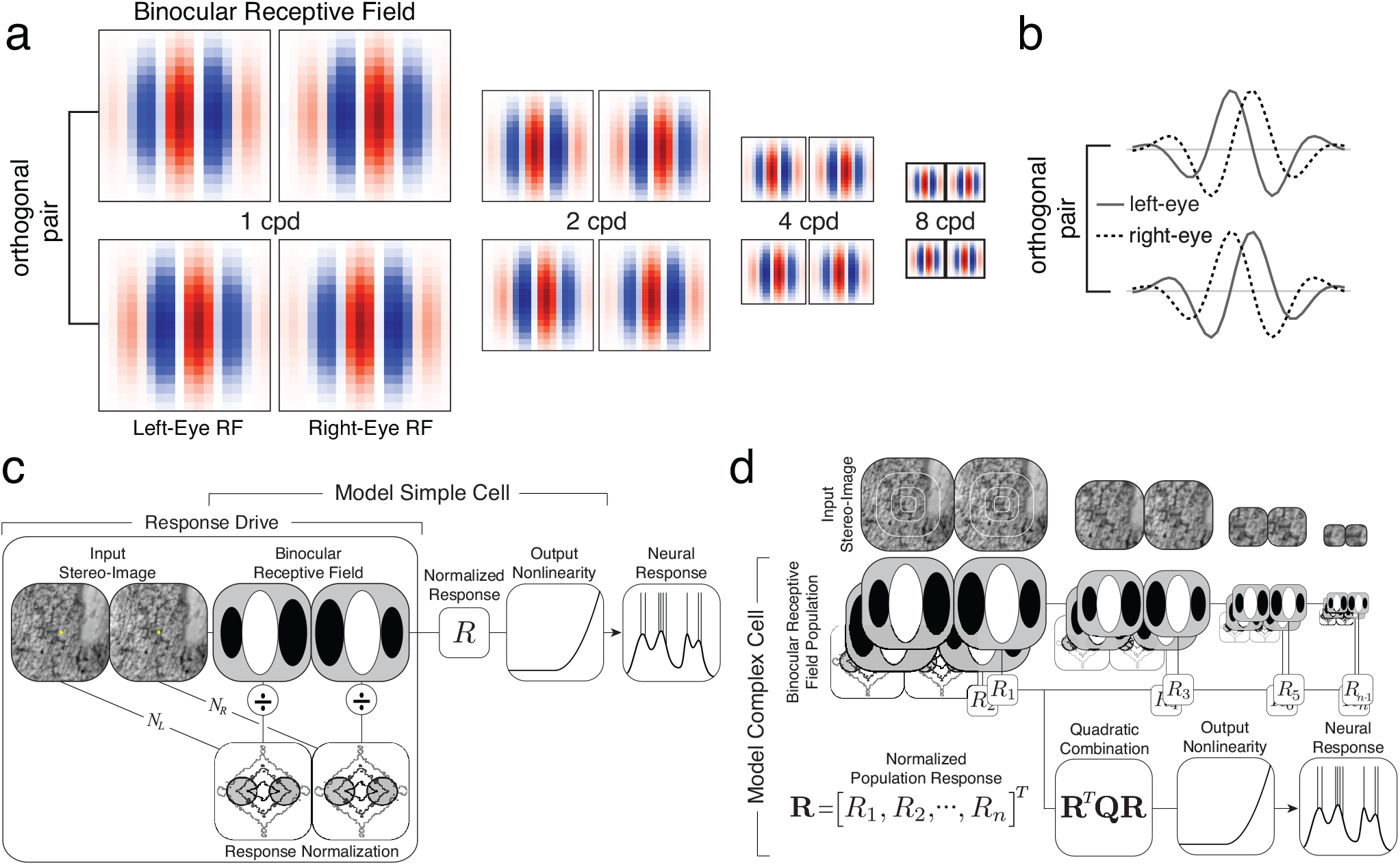
**(a)** Binocular receptive fields with preferred spatial frequencies of 1, 2, 4, and 8cpd. The left- and right-eye components of each binocular receptive field are phase shifted with respect to each other by 90º (−45º and +45º, or vice versa). **(b)** Receptive field profiles obtained by taking a horizontal slice through each of a binocular receptive field. As can be seen in (a), receptive fields (and profiles) are identical up to a spatial scale factor. **(c)** Binocular receptive fields are assumed to respond according to the following response model. The response is driven by a projection of the stimulus onto the receptive field modulated by (feature-specific) divisive normalization. (Note that normalizing the response after projecting the stimulus onto the receptive field, and normalizing the stimulus before projection onto the receptive field, yield computationally equivalent responses.) Finally, an output non-linearity, and a stochastic process that accounts for internal noise (not pictured), transform the normalized response field response into neural response. **(d)** Complex binocular neurons are modeled by the quadratic combination of responses from multiple differently-sized binocular receptive fields with different spatial frequency and phase preferences. Note that smaller receptive fields process a smaller region of the image. The current paper focuses its analysis on the statistics of the normalized receptive field responses— specifically, on the conditional probability of response *p*(**R**|*δ*) to natural stereo-image patches sharing the same fixation disparity. When the quadratic weights **Q** are appropriately chosen (see below), the neural response of the complex cell reports the likelihood that a stimulus with a particular disparity elicited the receptive field responses.

Neural responses of simple cells in primate early visual cortex are often modeled as arising from a set of processes: receptive-field-based filtering, divisive normalization, the addition of noise, and an output non-linearity which converts a response rate into spikes (Fig. 2c; Goris et al., 2024; Heeger, 1992; Albrecht & Geisler, 1991). Models of complex cell responses are more involved; before the output non-linearity, the responses of multiple receptive fields are quadratically combined (Fig. 2d).

Here, we analyze the distribution of normalized receptive field responses (i.e., response drives) rather than the noisy spiking responses of the neuron (see Fig. 2c), focusing on how the distribution of these responses *p*(**R**|*δ*) across many natural images with the same fixation disparity is affected by the form of normalization and by the spatial frequency preferences of the receptive fields. As will soon become clear, these response statistics entail that the normalized responses should be quadratically combined, consistent with standard descriptive models of complex cells (Fig. 2d).

### Narrowband vs. broadband normalization

#### Effects on response statistics

To examine how the responses of individual binocular receptive fields are driven by natural stimuli, we begin with stereo-images having zero fixation disparity, a viewing situation that arises when the eyes perfectly fixate a point on the surface of an object (see Fig. 1b). Then, we examine how the response statistics vary with fixation disparity and how well each receptive field by itself supports disparity discrimination.

The statistical properties of the responses to natural stereo-images depend heavily on the form of normalization. We consider two types of normalization: broadband normalization and narrowband normalization (Iyer & Burge, 2019). Broadband normalization is stimulus-specific, but feature-independent. It normalizes the receptive field response to each stimulus by all the stimulus contrast in the location of the receptive field; that is, every spatial frequency and orientation in the stimulus contributes equally to the normalization factor. Narrowband normalization is stimulus-specific and feature-dependent. It normalizes receptive field response by the stimulus contrast in the spatial frequency and orientation passband at the location of the receptive field. With narrowband normalization, stimulus features that are more similar to the preferred feature contribute more to the normalization factor.

The broadband-normalized responses of each individual binocular receptive field are approximately mean-zero Laplace-distributed (i.e., heavy-tailed); each response distribution has a small standard deviation and a kurtosis of approximately 6.0. By contrast, the narrowband-normalized responses are approximately mean-zero Gaussian-distributed; each response distribution has a larger standard deviation and a kurtosis of approximately 3.0 (Fig. 3ab). For both forms of normalization, these statistics are largely invariant to the preferred spatial frequency of the receptive field (Fig. 3c; also see Jaini & Burge, 2017 and Iyer & Burge, 2019). The kurtoses of the narrowband-normalized response distributions are also largely invariant to fixation disparity (Fig. 3d). The standard deviation of responses from each receptive field, however, varies periodically with fixation disparity, at a rate that depends on the preferred spatial frequency of the receptive fields (Fig. 4a).

**Figure 3:**
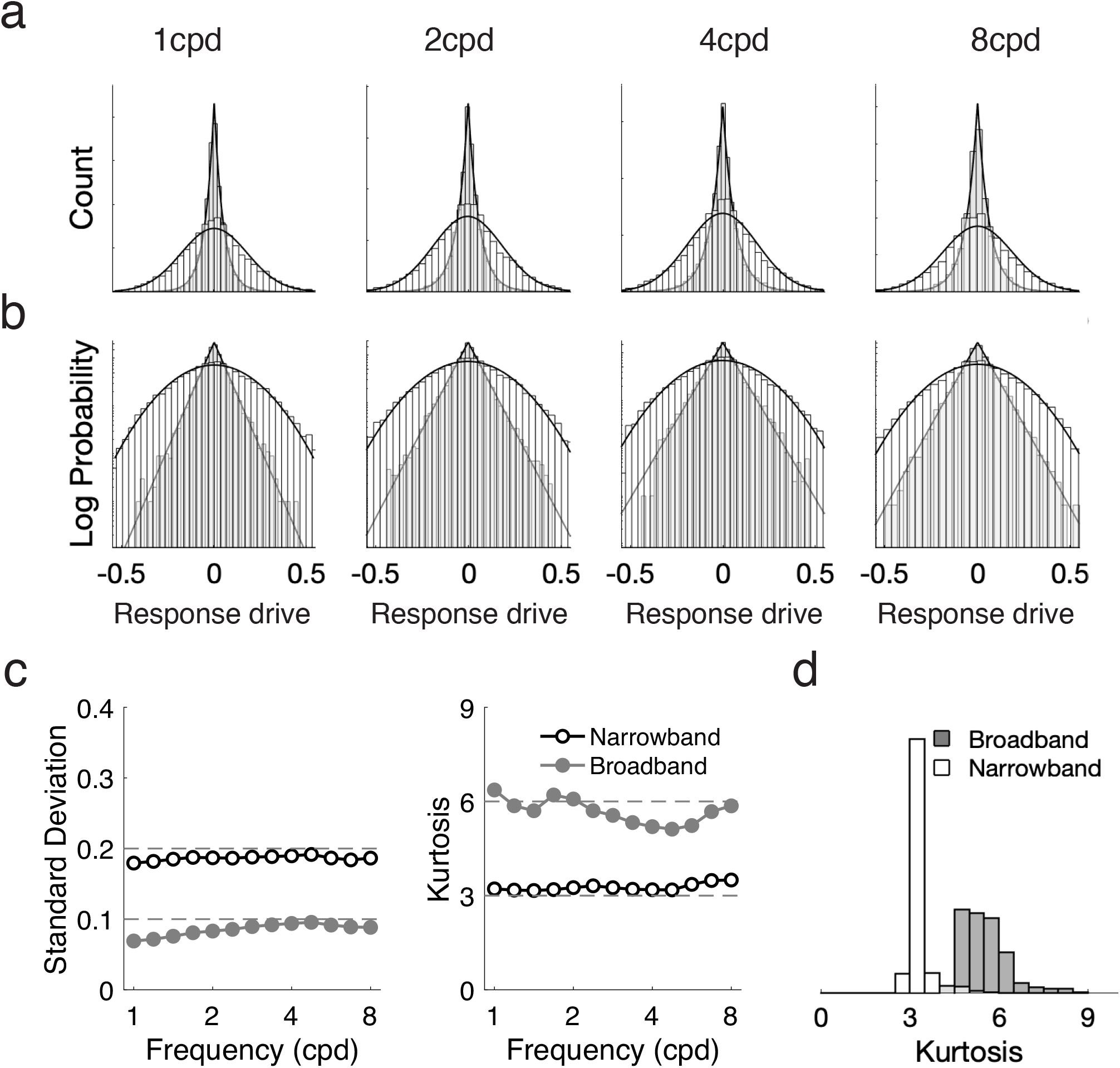
Binocular receptive field response statistics with broadband and narrowband normalization. **(a)** Distributions of broadband- and narrowband-normalized responses to natural stereo-images having a fixation disparity of 0 arcmin. The broadband- and narrowband- normalized responses are approximately Laplace and Gaussian distributed, respectively. **(b)** Same responses as in (a), but with log-probability transformed y-axis. **(c)** Response standard deviation (left) and response kurtosis (right) as a function of the receptive field’s preferred spatial frequency (see Fig. 2) for narrowband and broadband normalization (colors). Gaussian-distributed responses have a kurtosis of 3.0. Laplace-distributed responses have a kurtosis of 6.0. **(d)** Histogram of response kurtosis across all preferred spatial frequencies and fixation disparities.

**Figure 4:**
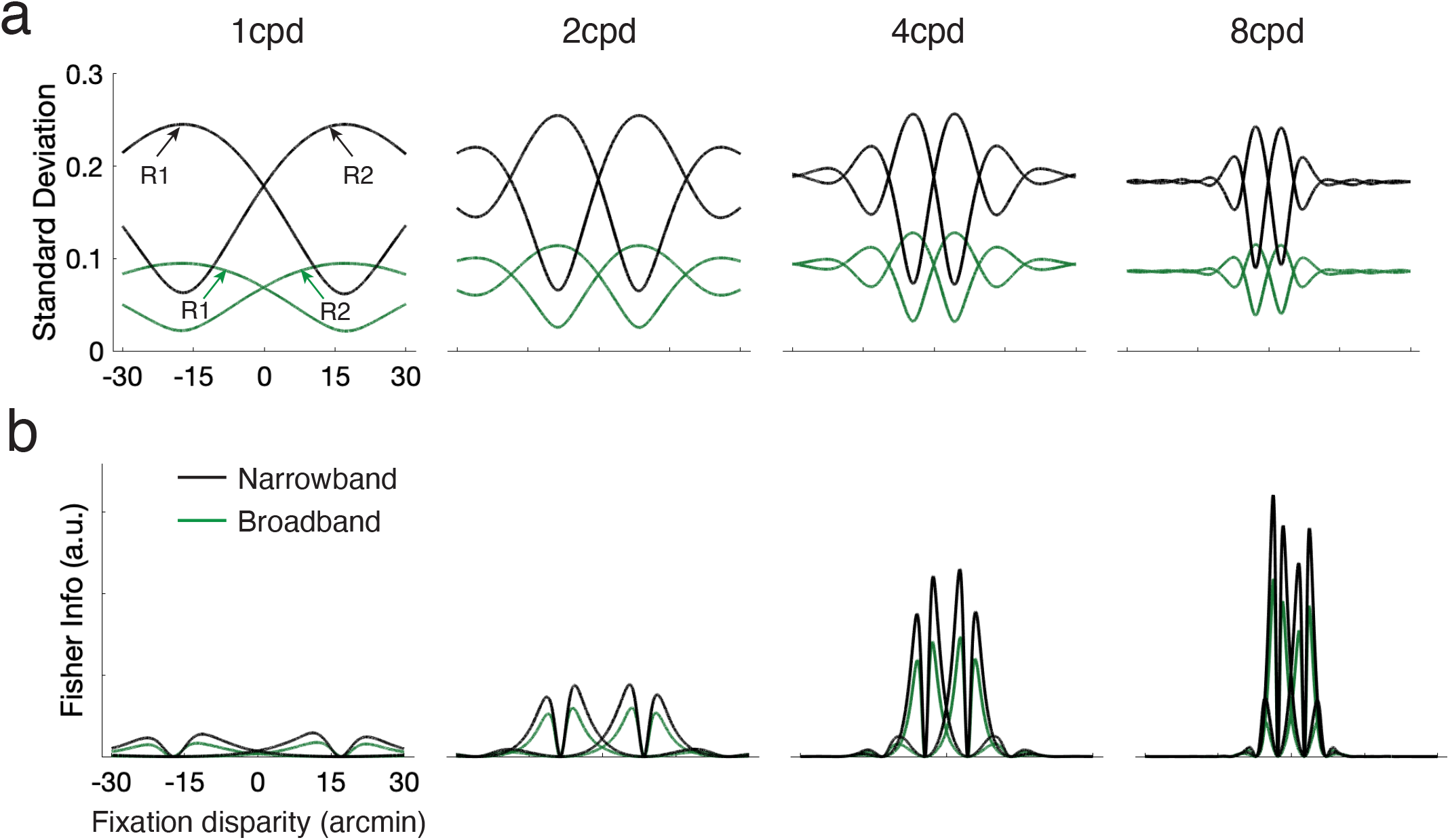
**(a)** Standard deviation of narrowband- and broadband-normalized responses (black and green, respectively) as a function of fixation disparity for individual receptive fields with different spatial frequency preferences (columns). **(b)** Corresponding Fisher information for each individual receptive field. Fisher information is consistently higher for narrowband- than for broadband-normalized responses.

### Narrowband vs. broadband normalization

#### Effects on encoding fidelity

To determine how well a latent variable—here, fixation disparity—can be discriminated from the responses of individual receptive fields, we compute Fisher information. Fisher information specifies the amount of information about a latent variable contained in a random variable—here, the receptive-field responses. Fisher information is equivalently given by the following two expressions

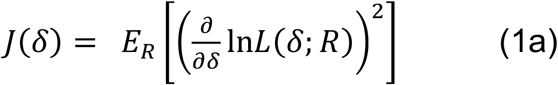

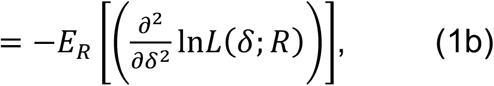

where *δ* is the fixation disparity, *R* is the receptive field response, and In*L*(*δ*; *R*) is the log-likelihood function over disparity, which can be computed by evaluating an observed response in the log of each response distribution *p*(*R*|*δ*). When the response distributions are mean-zero Gaussian, the response distributions are given by 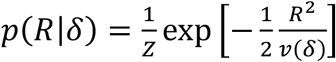 where *v*(*δ*) is the response variance at a particular disparity and *Z* is a constant. The log-likelihood that an observed response was elicited by a stimulus having a particular disparity is thus given by 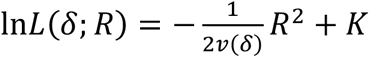 where *K =* −In*Z* is a constant.

Eq. 1a is the expected value of the squared derivative of the log-likelihood function, and Eq. 1b is the negative expected curvature of the log-likelihood function, at each fixation disparity. (Note that Eq. 1b holds only when the log-likelihood function is twice differentiable; Eq. 1a holds regardless.) In general, higher Fisher information implies lower discrimination thresholds (see below).

The Fisher information depends on the distributional form of the random variable, and on how the parameters of the distribution change with the latent variable. When the observed random variable is zero-mean Gaussian-distributed, as the narrowband-normalized responses tend to be at each disparity, Fisher information is given by

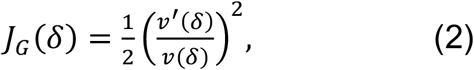

where *v*(*δ*) is the variance of the receptive field response at each disparity, and *v*′(*δ*) is the corresponding derivative with respect to disparity (see *Supplement* for derivation). When the observed random variable is zero-mean Laplace-distributed—as is the case with broadband normalized responses—the Fisher information is given by

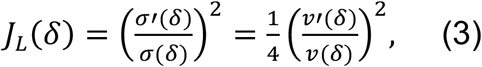

where *σ*(*δ*) is the standard deviation of the receptive field response at each disparity, and *σ*^′.^(*δ*) is the corresponding derivative with respect to disparity (see *Supplement* for derivation). The Fisher information can be equivalently expressed in terms of *v*(*δ*) the response variance at each disparity, and *v*^′^(*δ*) its corresponding derivative.

Clearly, the Fisher information in Gaussian-distributed responses is two times larger than that in the Laplace-distributed responses (see Eqs. 2 and 3), provided that the standard deviation (or variance) functions are the same, or are scale multiples of one another, as they approximately are here (see Fig. 4a). Narrowband-normalized responses thus contain more Fisher information than broadband-normalized responses, in each individual receptive field (Fig. 4b). So narrowband-normalization increases the fidelity with which fixation disparity is encoded. However, individual binocular receptive fields in isolation encode disparity rather poorly. Responses from pairs of binocular receptive fields contain substantially more Fisher information about fixation disparity, supporting better discrimination performance. Next, we analyze the response statistics of receptive field pairs, derive expressions for the Fisher information in responses from receptive field pairs and populations, and demonstrate that the benefit of narrowband normalization increases systematically with population size.

### Normalized responses from receptive-field pairs

#### Statistics and Fisher information

The joint narrowband-normalized responses *p*(***R***|*δ*) *= N*(*R*; **0, C**_*δ*_) to a collection of natural stereo-images having the same disparity are well-approximated as mean-zero multivariate Gaussian distributions (Fig. 5a). These response distributions are formally described by 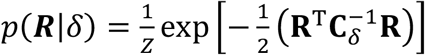 where 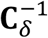 is the inverse covariance matrix associated with stereo-images having a particular disparity and *Z* is a constant that does not depend on the response. The log-likelihood that a stereo-image with a particular disparity elicited an observed population response is therefore given by

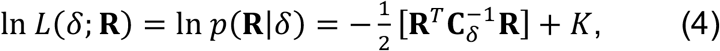

where *K =* −ln(*Z*) is an additive constant. Computing the log-likelihood (and likelihood) therefore requires quadratic combination of the receptive field responses. When the weights for quadratic combination are set equal to the inverse covariance matrix (i.e.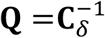 see Fig. 2d), neurons carrying out those computations would report the likelihood of that disparity. Such neurons would be perfectly suited for circuits implementing maximum likelihood—or Bayes-optimal—estimation in cortex (Burge & Geisler, 2014; Burge, 2020).

**Figure 5.**
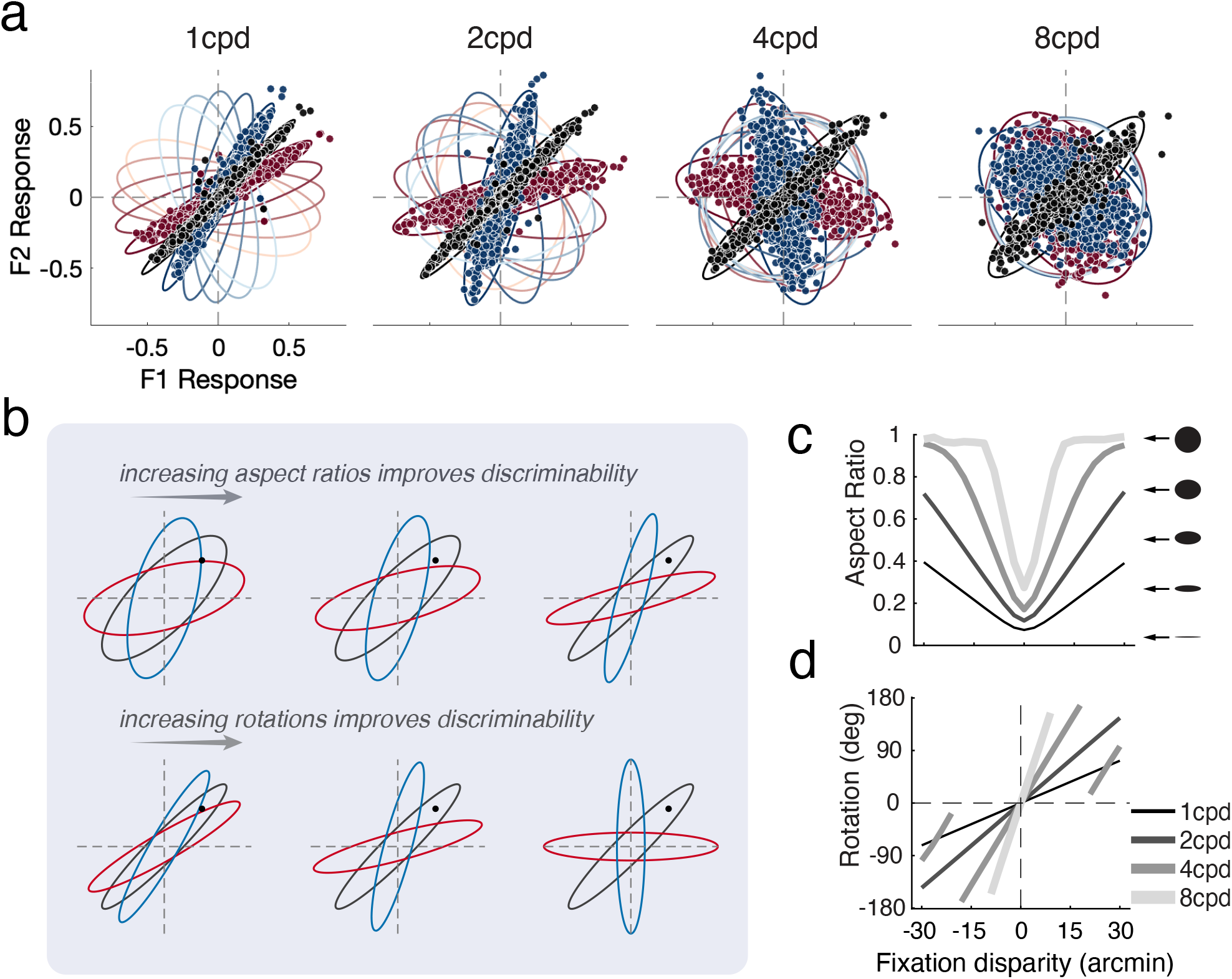
Joint receptive field response distributions. **(a)** Narrowband-normalized response distributions for the receptive field pairs defining each spatial frequency channel to a collection of images having different disparities (colors) Each point represents the response to a single stereo image. **b)** Schematic showing how changes in aspect ratio and in orientation of the response distributions for stimuli having uncrossed (blue), zero (black), and crossed (red) disparity, respectively. The black dot represents a joint response to a particular natural stereo-image patch associated with fixation disparity of 0 arcmin. **(c)** Aspect ratio and **(d)** rotation of each bi-variant normal distribution or “ellipse” shown in **a** as a function of fixation disparity. The rotation of an ellipse is defined as the difference between the orientation of its major axis and that of the ellipse associated with fixation disparity of 0 arcmin. The rotation of the ellipse associated with disparity of 0 arcmin is thus set to 0º.

The covariances of the response distributions clearly depend on both the fixation disparity and the spatial frequency preferred by the corresponding pair of receptive fields (Fig. 5). The manner in which the response covariances change with disparity suggests that pairs and populations of receptive fields should provide far more information about fixation disparity than individual receptive fields do (see below).

The covariance of each distribution can be characterized with the aspect ratio and orientation of each bi-variant joint distribution or ellipse (Fig. 5ab). This characterization is useful for examining how the distributions change with fixation disparity. (Note that the length of the principle axis of each ellipse, which corresponds to the principle eigenvalue, is essentially invariant to disparity, so aspect ratio and orientation fully capture the changes.) Increases in fixation disparity from zero cause the aspect ratios to approach 1.0 (i.e. cause the distributions to be more nearly circular; Fig. 5ac). Increasing the preferred spatial frequency (i.e. size) of the receptive fields causes the aspect ratios and the orientations to rotate more rapidly with fixation disparity (see Fig. 5cd).

Visual inspection of the response distributions suggests that different spatial frequencies should differentially support discrimination performance at varying fixation disparities; responses from distributions that overlap less are less confusable (Fig. 5). The Fisher information in the joint response distributions quantifies how this distributional overlap is related to discrimination thresholds. For responses from pairs (or populations) of receptive fields that are jointly distributed as zero-mean Gaussians, Fisher information is given by

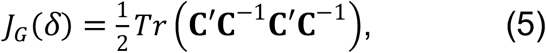

where *Tr*(.) is the trace operator and **C** is the covariance matrix (see *Supplement* for derivation; Mardia & Marshall, 1984). (In Eq. 5, the dependence of **C** on fixation disparity has been dropped for notational simplicity.) Because of how the response covariance changes with fixation disparity, there is a massive increase in Fisher information that a pair of receptive field responses provides over those provided by individual receptive fields in isolation (Fig. 6a). This finding confirms that pairs of binocular receptive fields should dramatically improve the encoding of disparity from natural stereo-images.

**Figure 6.**
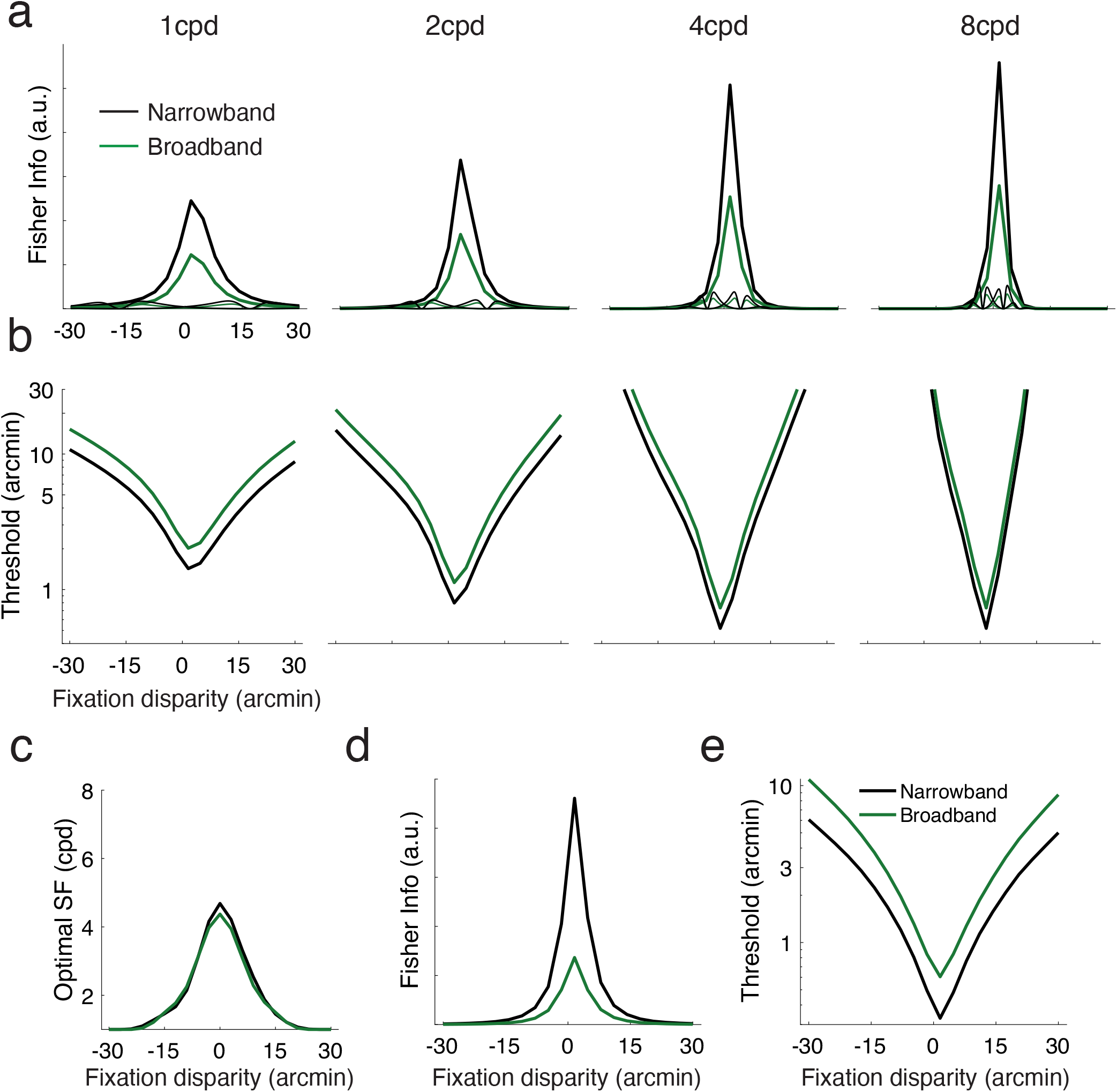
Functional population coding benefits afforded by narrowband normalization. **(a)** Fisher information in the joint narrowband-and broadband-normalized responses of each receptive field pair. Narrowband-normalized responses from each pair contain two times more Fisher information (see Eqs. 5 and 6). For reference, the Fisher information in the narrowband-normalized responses of each individual receptive field is also shown (replotted from Fig. 4b). Note the dramatic increase in information in the joint responses. **(b)** Disparity discrimination thresholds that are implied by the Fisher information in the joint responses. (**c**) Optimal spatial frequency as a function of fixation disparity. For each disparity, the optimal spatial frequency is the frequency preferred by the receptive field pair that gives rise to the highest encoding fidelity (i.e. Fisher information). (**d**) Fisher information in narrowband- and broadband--normalized responses in all four receptive field pairs (see *Method*s). (**e**) Corresponding disparity discrimination thresholds that are implied by the Fisher information.

The Fisher information in mean-zero multivariate Laplace-distributed responses is given, to close approximation, by

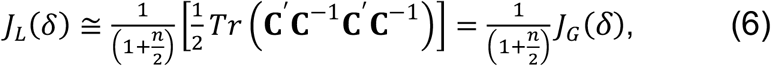

where *n* is the number of dimensions (see *Supplement* for derivation). The equation implies that, for a receptive field pair (i.e. *n*=2) composing a spatial frequency channel, there is 2x less Fisher information in bivariate broadband-normalized (Laplace-distributed) responses than in narrowband-normalized (Gaussian-distributed) responses (Fig. 6b), provided that the covariances are equal up to a scale factor, as they approximately are here (see Fig. 4a).

Eqs. 5 and 6 further imply that narrowband normalization provides an increasingly large advantage over broadband normalization as the number of receptive fields (dimensions) supporting discrimination increases; the Fisher information is (1 + *n*/2) times larger. Given that complex cells in cortex, including disparity-selective cells, tend to be driven by many more than two (i.e. ten or more) subunit receptive fields (Fig. 2d) (Rust et al., 2005; Tanabe, Haefner, Cumming, 2011), the functional coding advantage afforded by narrowband over broadband normalization in neural systems should be substantial.

The reciprocal of Fisher information gives the Cramer-Rao lower bound on the variance of an unbiased estimator, which directly predicts the discrimination thresholds that can be measured in a psychophysical experiment. However, unbiased estimators are not always realizable and estimators are often biased. Fortunately, it can be shown that Fisher information specifies the smallest possible discrimination thresholds via an expression that is invariant to whether the estimator is biased or not (see *Supplement*). Specifically,

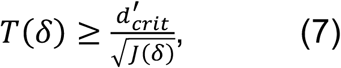

where *T*(*δ*) is the discrimination threshold as a function of fixation disparity, and 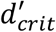 is the criterion *d*-prime. (For simplicity, but without loss of generality, we assume that the criterion d-prime is 1.0, corresponding to 76% correct threshold in a two-interval two-alternative forced choice task.) Discrimination thresholds implied by the Fisher information (see Eq. 7) in responses of receptive field pairs are square-root-of-two times smaller for the (Gaussian-distributed) narrowband-normalized responses than for the (Laplace-distributed) broad-band responses (Fig. 6b).

Moreover, we note several findings that are invariant to the type of normalization. First, small fixation disparities are encoded with greater fidelity than large disparities, a result that does not depend on the preferred spatial frequency of the binocular receptive fields. Second, binocular receptive fields preferring lower spatial frequencies code disparities over a larger range than those preferring higher spatial frequencies. Third, binocular receptive fields that prefer higher spatial frequencies are better at encoding small disparities, and those that prefer low spatial frequencies are better at encoding large disparities (Fig.6c). All of these findings have analogs in the psychophysical and neurophysiological literatures (Blakemore, 1970; DeAngelis & Cumming, 2001; McKee, Levi, Bowne, 1990; Prince et al., 2002; Schor & Wood, 1983; Siderov & Harwerth, 1993; Smallman & MacLeod, 1994; see *Discussion*).

### Population coding

Perceptual systems should use all relevant information to perform critical tasks. Thus, the information provided by all receptive fields in a population, spanning various spatial frequency preferences, should be used in concert. To examine how encoding fidelity improves when the population response is used, we compute the Fisher information from the normalized responses receptive field pairs that fully tile the 1-8cpd range with different densities. Encoding fidelity improves—Fisher information increases (Fig. 7a), and discrimination thresholds decrease commensurately (Fig. 6de and Fig. 7b)—for all fixation disparities, as compared with any spatial frequency channel in isolation. The improvement in encoding fidelity and discrimination performance increases monotonically, but with decreasing returns, as the density of receptive fields (i.e., number of spatial frequency channels) tiling the spatial frequency range increases (Fig. 7cd).

**Figure 7.**
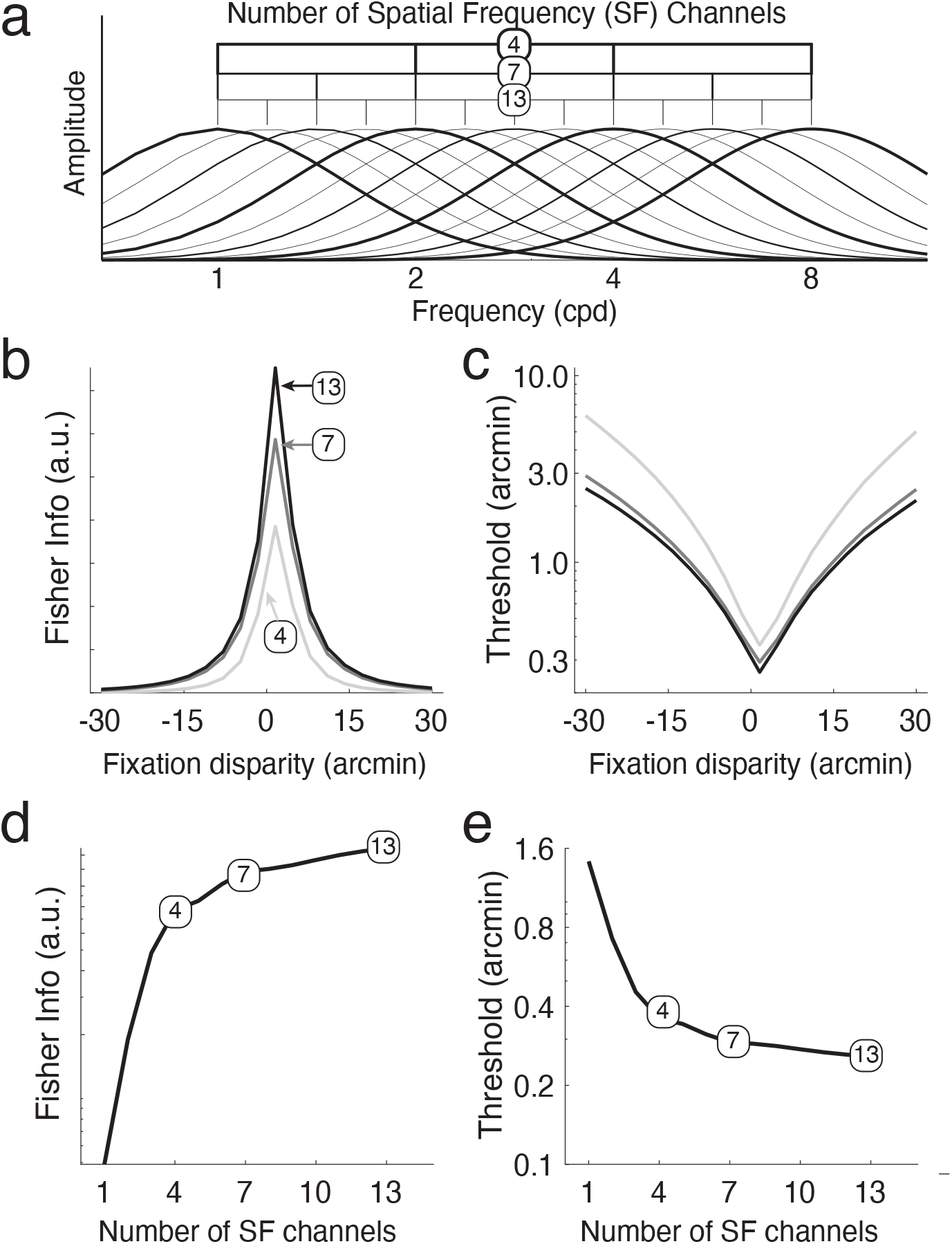
Functional benefits of population coding. **(a)** Depiction of the density with which different numbers of spatial frequency (SF) channels (4, 7, or 13) tile the spatial frequency range 1-8cpd. **(b)** Fisher information in narrowband-normalized responses for receptive field populations associated with different numbers of channels as a function of fixation disparity. Each curve shows Fisher information for a different number of receptive field pairs tiling the 1-8cpd spatial frequency range (colors). Note the diminishing returns as the density of tiling increases. **(c)** Disparity discrimination thresholds that are implied by the Fisher information in (b). (**d**) Fisher information and **(e)** discrimination (i.e. detection) thresholds at fixation disparity of 0 arcmin as a function of the number of spatial frequency channels.

The lowest disparity discrimination threshold (i.e. the disparity detection threshold) is a fraction of an arcmin, and discrimination thresholds rise exponentially with fixation disparity from the detection threshold (see Burge & Geisler, 2014). These computational results, and others, are in close alignment with classic findings in the psychophysical literature (see *Discussion*).

The fact that Fisher information increases, and discrimination thresholds decrease when receptive field responses are appropriately combined across spatial frequency channels is expected: each channel is useful, so combining useful information should improve performance. However, the statistical approach used here reveals at least one non-obvious benefit: the presence of multiple channels allows the system to adaptively adjust to the variable usefulness of each channel for individual stimuli, not just on average. That is, for a given stimulus, when one channel provides little or no useful information about fixation disparity, other channels can compensate, ensuring reliable performance across different conditions.

Consider stimuli that elicit receptive field responses in the confusion zone of a low spatial-frequency channel, where the response magnitude ‖***R***_*f*_‖ nearly equals zero (Fig. 8a, 1cpd). The disparity associated with those stimuli will be hard to discriminate because of the heavy overlap of the response distributions near zero. Those same stimuli, however, will elicit differential responses in the next higher spatial-frequency channel. While some of these new responses will still fall in the confusion zone of the new channel, others will be large and outside the confusion zone, thereby enhancing discrimination performance (Fig. 8a, 2cpd). In the next frequency channel, some stimuli that produce responses in the confusion zones of the first two channels will generate large responses in the third channel, enabling good performance on the basis of the third channel (Fig. 8a, 4cpd), and so on (Fig. 8a, 8cpd). The same principle can hold in the other direction (Fig. 8b).

**Figure 8:**
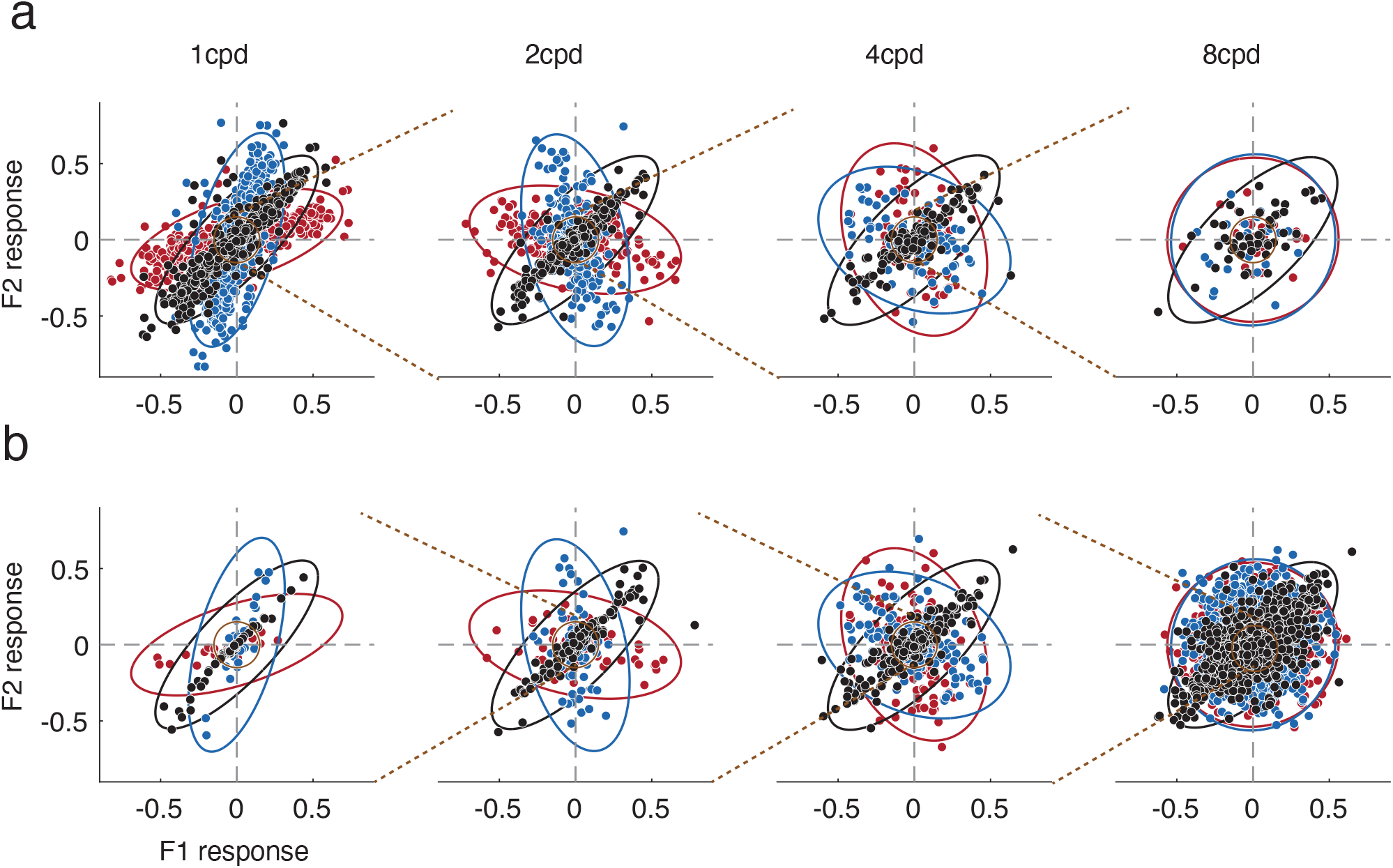
Joint response distributions of binocular receptive field pairs having preferred spatial frequencies of 1, 2, 4, and 8 cyc/deg to natural stereo-images with uncrossed (blue), zero (black), and crossed (red) disparity. When stimuli elicit joint responses in the “confusion zone” (orange circle; ‖**R**_*f*_‖ ≤ *d* where *d* is the distance from the origin) of a given frequency channel, the associated fixation disparities are harder to discriminate than when responses are farther away. Stimuli that cause responses within the confusion zone of one frequency channel can elicit more discriminable responses in another. **(a)** Stimuli that cause responses within the confusion zone for a low spatial frequency channel (e.g., 1 cyc/deg) cannot be discriminated based on those responses alone. But a substantial subset of those stimuli elicits responses outside of the confusion zone of a higher spatial frequency channel (e.g. 2cyc/deg). Furthermore, stimuli that cause responses in the confusion zones of the first two spatial frequency channels can elicit responses outside the confusion zone of an even higher spatial frequency channel (e.g. 4cyc/deg), and so on. This complementary processing across channels enables good overall discrimination performance. **(b)** The same pattern holds when the initial confusion zone is for a binocular receptive field pair that prefers a high spatial frequency (e.g. 8cyc/deg), and the disambiguating channels are lower frequency.

For some fixation disparities, certain spatial frequency channels are more useful than others on average across stimuli (see Fig. 5). However, with a particular stimulus, Fig. 8 shows that the usefulness of different spatial frequency channels depends not just on which channels are most useful on average, but on the specific constellation of features that defines each stimulus and on how the features cause the within-channel receptive fields respond. That is, for given stimulus, the quality of the information in a certain spatial frequency channel is signaled by the magnitude of the within-channel receptive-field responses. A stimulus that produces receptive field responses in the confusion zone of the 1cpd channel, for example, indicates that the channel provides no relevant information for latent variable discrimination (see Fig. 8a). Under such circumstances, the channel should be ignored. The rules for computing the likelihood do just this.

Despite the fact that the rules for computing the likelihood from each stimulus are deterministic (i.e. they are fixed, or passive, in that the same rules are used to process each stimulus), they automatically and adaptively prioritize the information that each channel provides on a stimulus-by-stimulus basis, according to the quality of the within-channel information. Specifically, the magnitude of the within-channel receptive field responses signals the usefulness of each channel.

To sum up, the statistics of receptive field response dictate the deterministic rules of computation for computing the likelihood; they depend on the quadratic combination of responses (see Fig. 2d) with weights 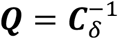 (see Eq. 4). The stimulus-by-stimulus responses determine the influence that each feature in a particular stimulus should have in computing the likelihood. Hence, with a set of fixed (possibly synaptically-implemented) weights, the magnitude of receptive field responses in each channel automatically determines the influence each channel should have on the computation of the likelihood, and of the eventual estimate, for each individual stimulus.

## Methods

### Natural stereo-image dataset

The dataset used for the analysis in this paper consisted of 91 stereo-images (1920×1080 pixels) of natural scenes taken on and around the University of Texas at Austin (Burge et al., 2016). The left- and right-eye images were photographed from one human interocular distance (i.e. 65mm) apart, and were each co-registered pixel-wise with laser-based distance measurements having millimeter-scale precision. The nodal points of the photographic camera and the laser-range scanner were aligned with a custom-built robotic gantry such that there were no half-occlusions between the photographic images and distance “images” of each scene. The field of view of each image subtended 35.8ºx21.1º of visual angle.

In a binocular visual system, surface points in the scene tend to be imaged in each image at different locations because each eye has a different vantage point on the scene. Such local differences in the positions of these corresponding points are binocular disparities. Biological visual systems must estimate which points in the left-eye image correspond to which points in the right-eye image from image data. Solving this correspondence problem is a core computational challenge underlying stereo-depth perception.

Fortunately, because of the properties of the dataset, ground-truth corresponding points can be computed directly from the distance data with arc second precision. Custom software, which is based on the principles of ray-tracing, is available online (Iyer & Burge, 2018; https://github.com/burgelab/StereoImageSampling). Each pair of left- and right-eye corresponding points depicts a particular point on a surface in the three-dimensional scene (see Fig. 1a).

Stereo-image patches were sampled with fixation disparity by simulating a virtual human observer that converges his/her eyes in front, on, or behind a surface point in the 3D scene along the cyclopean line of sight. These eye postures cause the surface point to have uncrossed, zero, and crossed fixation disparities, respectively (Fig. 1b). Fixation disparity is rigorously given by the difference in vergence angle (i.e. the angle between the left- and right-eye lines of sight) between the currently fixated location and the vergence demand of the surface point in question

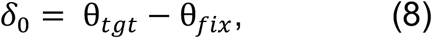

where θ_*tgt*_ is the vergence angle demanded by the target point on the surface and θ_*fix*_ is the vergence angle associated with the fixated location. Clearly, if the eyes are perfectly fixated on the surface point, the surface point is imaged with zero fixation disparity.

Natural scenes are composed of objects that have three-dimensional structure, so images of natural scenes tend to depict scenes that vary in depth. The amount of local depth variability can vary significantly across image patches. We index the amount of local depth variability in a patch with disparity contrast. Disparity contrast is given by

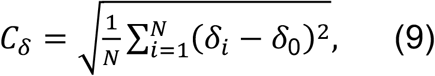

where *δ*_*i*_ is the disparity at each pixel *i, δ*_0_ is the fixation disparity at the central pixel (i.e. the corresponding point), and *N* is the number of pixels within the image. (For a given fixation distance, disparity contrast perfectly covaries with local depth variation.) Natural image patches with low and high disparity contrast are shown in Fig. 1c. Patches with low disparity contrast (< 2arcmin)—which are the focus of the current article—occur far more often than patches with high disparity contrast in natural scenes (*Supplementary* Fig. S2; see also Iyer & Burge, 2018).

### Response model

#### Response model: Binocular receptive fields

Binocular receptive fields have a left-eye and a right-eye component. Each eye’s component is modeled as a vertically-oriented Gabor, a vertical sine wave multiplied by a Gaussian envelope. The spatial frequency octave bandwidth was 1.2 and with an orientation bandwidth of 42º. The left-eye component of each binocular receptive field had a phase-shift of either -45º or +45º, and the right-eye component had a phase-shift of either +45º or -45º, such that every binocular receptive field had a preferred phase disparity of either -90º or 90º. The preferred fixation disparity, in degrees of visual angle, is given by

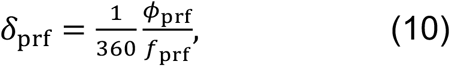

where *Φ*_prf_ is the preferred phase disparity in degrees and *f*_prf_ is the preferred spatial frequency.

Each model binocular receptive field preferred one of thirteen spatial frequencies equally-spaced in the log-domain between 1 and 8 cyc/deg. Fig. 2a shows four pairs of binocular receptive field pairs, which are in quadrature phase, with spatial frequency preferences of 1, 2, 4, and 8 cyc/deg. Fig. 2b shows horizontal slices through the binocular receptive field that selects for the lowest spatial frequency.

The responses of model binocular receptive fields (see below) to natural stereo images were computed for 121 fixation disparities between -30 to 30 arcmin, in 0.5 arcmin steps. For each disparity level, we sampled a total of 21784 binocular image patches from 91 natural stereo-images. The spatial region of the image patch that contributes to each normalization factor is matched to the size of the receptive field. The largest receptive field, which prefers 1 cyc/deg, was 72×72 pixels in size. The smallest receptive field, which prefers 8 cyc/deg, was 9×9 pixels in size.

#### Response Model: Divisive normalization

A binocular receptive field receives visual input from both the left and right eyes. In general, the left-and right-eye images will tend to be slightly different, due to each eye’s different vantage point on the depth-varying scene. The response of a binocular receptive field is given by the sum of the normalized outputs of the left- and right-eye receptive field components: *R*_L_ and *R*_R_. Specifically, the response of a binocular receptive field is given by

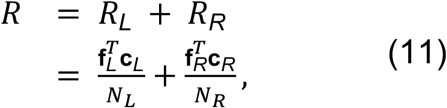

where **f**_*L*_ and **f**_*R*_ are, respectively, the left-eye and right-eye linear receptive fields, **c**_*L*_ and **c**_*R*_ are the Weber contrast images for the left and right eye, and *N*_L_ and *N*_R_ denote the normalization factors for the linear responses of the left- and right-eye receptive field components.

Two types of normalization are considered: broadband and narrowband. Broadband normalization factors for the left- and right-eye receptive fields are given by

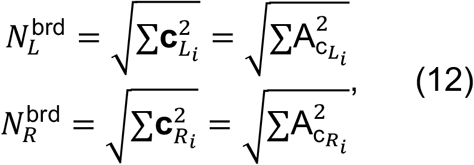

where 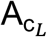 and 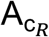 are the amplitude spectrum of the left- and right-eye contrast images, respectively. Parseval’s theorem states that the total energy of a contrast stimulus equals the total power of the (amplitude) spectrum. Thus, Eq.12 shows that all stimulus features of the binocular image contribute equally to the factor of broadband normalization. (Note that the normalization factor could alternatively be computed using a method that computes a single binocular normalization factor across the left- and right-images (e.g., Hou, Nicholas, & Verghese, 2020). However, the binocular receptive field responses are largely invariant to this subtlety (see Fig.S1 in *Supplement*)

The narrowband—or feature-specific—normalization factors for the left- and right-eye receptive fields are given by

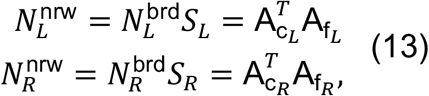

where *S*_*L*_ is the cosine similarity in the amplitude spectrum between the left-eye image and the left-eye receptive field and *S*_*R*_ is the same quantity computed for the other eye. These quantities provide a phase-invariant measure of similarity between the images and receptive fields in the two eyes and are specifically given by

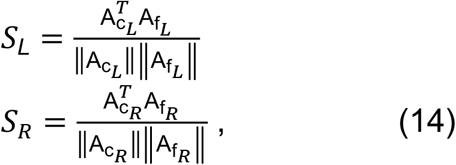

where the ‖.‖ operator indicates the L2 norm.

Eq.14 entails that different features of the stimulus contribute unequally to the narrowband normalization factor. Stimulus features that are similar to the preferred features of the receptive field contribute more than those that are dissimilar. When the left- and right-eye input perfectly matches its receptive field (i.e. *S*_*L*_ = *S*_*R*_ *=* 1), the narrowband and broadband normalization factors are identical. In most cases, however, *S*_*L*_ and *S*_*R*_ are smaller than 1.0, which causes the narrowband normalization factor to be smaller than the broadband normalization factor and a relatively larger response.

## Discussion

In this study, we demonstrated the functional value of narrowband (or feature-specific) response normalization in the neural coding of binocular disparity—a powerful cue to depth—in stereo-images of natural scenes. Using a database of natural stereo-images and a computational approach, we examined how common components of neural response models affect the statistical properties of binocular receptive field response. The analysis of response statistics provides insights into the design principles that should underlie population coding in image-computable models of natural tasks. Specifically, by quantifying the Fisher information about binocular disparity in the receptive field responses to natural images, we demonstrate how the spatial frequency preferences of the encoding receptive fields shape the response statistics, dictate the optimal combination rules across spatial frequencies, and determine the precision with which different binocular disparities are discriminated. The predicted patterns of discrimination performance are in line with classic and recent reports in the literature. The current approach represents a promising method for determining how neural systems should be designed and how patterns of human performance can be predicted by the task-relevant statistical properties of natural images and scenes.

### Functional benefits of feature-specific normalization

Contrast normalization is a ubiquitous neural computation. It was initially introduced approximately thirty years ago to help account for the contrast response function of cells in early visual cortex (Geisler & Albrecht, 1991; Heeger, 1992). In the three decades since, it has been invoked to account for many other properties of neural response (Carandini & Heeger, 2012). In recent years, feature-specific (or narrowband) normalization has been shown to improve accounts of neural responses to natural images (Burg et al., 2021; Coen-Cagli et al., 2012; Coen-Cagli et al., 2015; Fang et al., 2023; Goris et al., 2024), and to improve latent-variable encoding-decoding from natural images (Burge & Geisler, 2014; Jaini & Burge, 2017; Iyer & Burge, 2019).

Here, we derived analytic expressions and performed empirical analyses that quantify the computational benefit of narrowband over broadband normalization when the task is to discriminate binocular disparity from natural stereo-images. This improvement occurs because narrowband response normalization formats the receptive field responses to natural images so that they are approximately Gaussian-distributed (also see Wainwright & Simoncelli, 2000; Schwartz & Simoncelli, 2001; Iyer & Burge, 2019), and can be more readily decoded. This increase in Fisher information corresponds to a decrease in discrimination threshold by the square root of the same factor. So narrowband normalization should be favored by evolutionary processes that seek to maximize an organism’s ability to precisely discriminate the depth structure of natural scenes, especially when the relevant neural populations are large.

### Receptive field responses, optimal computations, and a generalized energy model

The underlying presumption of this manuscript is that the normalized receptive field response is the stage of neural processing best suited to a computational analysis of the information in natural images supporting stereo-depth discrimination (Marr, 1982). It is widely appreciated that divisive normalization helps account for the response properties of simple and complex cells in cortex (Albrecht & Geisler, 1991; Heeger, 1992; Carandini, Heeger, Movshon, 1997; Carandini & Heeger 2012). But spiking neural responses are quite obviously not Gaussian-distributed (Baddeley et al., 1997). To what degree does the current findings relate to real neurons in cortex and to mainstream approaches in computational neuroscience for analyzing their response properties?

Analyzing receptive field response statistics rather than spiking responses departs from practices in some areas of computational neuroscience (e.g., Hubel & Wiesel, 1962; Shadlen & Newsome, 1998; Rieke et al., 1999), but it aligns with practices in others. Subunit-based methods for neural systems identification—for example, spike triggered covariance analysis (Cook & Forzani, 2009; Rust et al., 2005), the generalized quadratic model (Park et al., 2013), and other related approaches (McFarland, Cui, Butts, 2013)— assume that quadratic combination of subunit receptive field responses underlies the intracellular voltage and spiking responses of complex cells. Such computations extract all information in the subunit receptive field population response up to second order, and do so optimally when the subunit responses are Gaussian-distributed (Jaini & Burge, 2017). So understanding the computations assumed by popular models in the context of Gaussian-distributed subunit responses links popular descriptive models to a normative account that is grounded in the statistics of natural scenes.

Given the mean-zero multivariate Gaussian-distribution of population response, computing the log-likelihood that a stereo-image with a particular disparity elicited an observed response therefore requires quadratic combination of the receptive field responses (see Eq. 4). Real neural systems that carry out such computations, or close approximations to them, would be well-suited to implementing maximum likelihood or Bayesian estimators. Also, quadratic combination is a hallmark of the disparity energy model, although the optimal computations—here, with weights given by the (inverse) covariance matrix 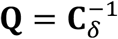—are best-described as a generalized version of it (Jaini & Burge, 2017; Herrera & Burge, 2024). The current results thus provide a normative justification for a classic descriptive model of neural response. Analyzing how natural stimuli stimulate sensors that are representative of those in real biological systems can provide new insight into why neural systems compute as they do.

### Combining information across spatial frequency channels

Previous computational reports have shown that multiple different spatial frequencies are differentially useful for coding binocular disparity (Smallman & Macleod, 1994; Read & Cumming, 2007; Qian & Zhu, 1997). But these studies employ heuristics for how to combine information across spatial frequency channels. The current analysis provides a principled approach for determining how information across multiple spatial frequency channels should be combined (Burge & Geisler, 2014). Specifically, it specifies the weights for how information should be quadratically combined across receptive field responses—and hence across spatial frequency channels—to compute the likelihood of a given disparity, which are dictated by the covariance of responses across stimuli having that disparity (see Eq.15). Additionally, it provides closed form expressions for how that information determines discrimination thresholds, arguably the most common psychophysical measure for assessing sensory-perceptual performance.

The fact that the receptive field responses—within and across spatial frequency channels—to natural stimuli associated with a particular fixation disparity are well-approximated as multivariate mean-zero Gaussian dictates that those responses should be quadratically combined to obtain the likelihood of each disparity. The covariance of those response distributions specifies the weights with which that quadratic combination should occur, and therefore how information from different spatial frequencies should be combined (see Results).

The general approach of examining the statistical properties of receptive field responses to different natural stimuli that are associated with the same value of the latent variable should be useful for developing normative models of many sensory-perceptual tasks, within and outside of vision (Burge, 2020). Such models will be important for gaining a principled understanding as to why neurons carry out the computations that they do.

### Gabor vs. task-optimized receptive fields

The current analysis analyzed the response of biologically-plausible Gabor-shaped binocular receptive fields and used Fisher information to compute the theoretical lower bounds on disparity discrimination thresholds implied by the statistical properties of receptive field response. Because the receptive fields used here were not themselves optimized for the task of disparity-discrimination with natural stimuli (Jaini & Burge, 2017), it cannot be claimed, that a system making use of task-optimized receptive fields could not exceed the indicated performance limits. Previous attempts to learn, from natural images, receptive fields optimized for disparity estimation have yielded Gabor-like receptive fields (Burge & Geisler, 2014; Burge & Jaini, 2017; Jaini & Burge, 2017) and performance patterns similar to those reported here. So the current results should therefore be considered representative. Also, the results suggest that feature-specific normalization should be incorporated into routines for learning task-optimized receptive fields, for the simple reason that it formats their responses so that the task-relevant latent variable can be more precisely decoded.

### Stimulus variability and stereo-depth discrimination

A central problem faced by the visual system is to precisely discriminate and accurately estimate behaviorally-relevant latent variables in the face of substantial natural stimulus variability. The difficulty is that there are an uncountable number of natural stereo-images that can share a given fixation disparity. This external stimulus variability inevitably limits the precision of perceptual estimation (or categorization). One way to formulate the goal of sensory-perceptual processing is to determine the computations that maximize accuracy while minimizing the deleterious effect of natural nuisance variability to the maximum possible extent (Burge & Jaini, 2017; Chin & Burge, 2020).

The current analysis examined how disparity discrimination performance is limited by natural image variation associated with nearly flat surfaces—i.e. variation across stereo-images with near-zero disparity contrast (see *Methods*). Although flat surfaces—or, more precisely, stereo-images with low disparity contrast—dominate the visual diet (see Fig. S2), it is not uncommon for a patch of image to depict a surface that is not flat, or to depict two surfaces at different depths (Iyer & Burge, 2018). How such local depth variability impacts disparity estimation and discrimination in natural scenes is a topic of ongoing research.

### Predicting human psychophysical data

The current computational results predict many well-established aspects of human disparity discrimination performance in the literature. Human disparity discrimination thresholds tend to rise approximately exponentially with fixation disparity (Badcock & Schor, 1985; Blakemore, 1970; McKee, Levi, & Bowne, 1990), and the lowest disparity discrimination threshold (i.e. the disparity detection threshold) is approximately 30 arcsec under typical (non-vernier) conditions (Blakemore, 1970; McKee, Levi, & Bowne, 1990). Under ideal (vernier) conditions, the detection threshold can reach 5 arcsec, approximately one sixth the width of a foveal cone photoreceptor (Stevenson, Cormack, Schor, 1989). However, the experiments that established these findings were all carried out with artificial stimuli: gratings (Badcock & Schor, 1985), random dot stereograms (Stevenson et al., 1989; McKee et al., 1990), and individual dichoptically-presented dots (Blakemore, 1970).

Recent findings show that similar patterns of performance occur when human observers are tasked with discriminating depth from natural stereo-images. White & Burge (submitted) tasked human observers with discriminating depth from small (∼1deg) patches of stereo-images that depicted surfaces in natural scenes. Some of the surfaces were nearly flat. Other surfaces were substantially bumpier. For both surface types, discrimination thresholds increased exponentially with fixation disparity. Thresholds associated with bumpier surfaces were larger than those with flat surfaces by a multiplicative scalar. The current article analyzed only surfaces that were nearly flat (see Fig. 1). A preliminary analysis on bumpy surfaces, using the same dataset and computational framework used in the rest of the paper (Ni & Burge, 2023), predicts multiplicative threshold increases that are similar to those that have been observed psychophysically. As mentioned above, a more detailed analysis is in progress.

## Conclusion

Computational-level investigations of how properties of natural stimuli and components of neural response models alter the form and fidelity of the encoded information can help one evaluate the design of real neural systems, discover the functional role of different components, and understand how perceptual performance in natural viewing is supported by underlying neural activity. The task and the stimulus ensemble determine the level of nuisance variability. The receptive field determines the information that is extracted, or selected, from each stimulus. And the form of normalization determines how the selected information will be formatted. The current analysis demonstrates a functional advantage for feature-specific (i.e. narrowband) normalization, quantifies the relative usefulness of different spatial frequencies in disparity discrimination, and predicts human performance in stereo-depth perception tasks.

## Acknowledgements

This work was supported by NIH Grant R01-EY028571 to JB from the National Eye Institute and the Office of Behavioral and Social Sciences Research, and by discretionary funds from the University of Pennsylvania. We thank Raymond Kan for helpful discussion about expectations of quadratic forms in elliptical random variables.

## Supplement

### Fisher information in mean-zero Gaussian-distributed Responses

The zero-mean Gaussian distribution is given by

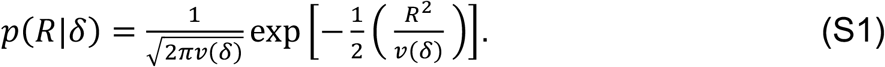

For receptive field responses *R* to be useful for discriminating a latent variable—here, fixation disparity—the conditional response distributions *p*(*R*|*δ*) must change with that latent variable. For a zero-mean Gaussian distribution, this requirement entails that variance *v*(*δ*) is a function of the latent variable. The expression for the Fisher information that is contained in such Gaussian-distributed responses is obtained by plugging the expression for the Gaussian probability density (Eq. S1 into Eq. 1b in the main text. Specifically,

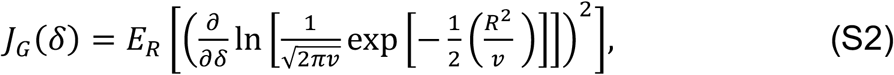

where, for notational simplicity, the dependence of the variance on the latent variable has been dropped.

Using the product rule for logarithms and distributing the derivative operator

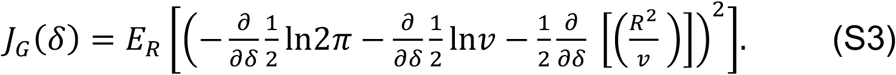

Taking the derivatives

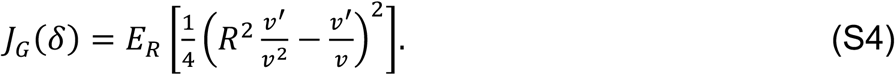

Expanding the square

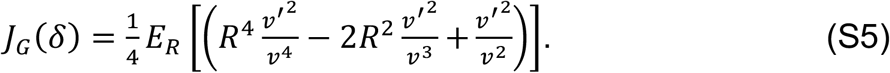

If *R* is mean-zero and Gaussian distributed, *E*[*R*^2^] *= v* and *E*[*R*^4^] *=* 3*v*^2^, so application of the expectation operator gives

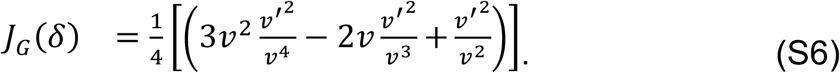

Canceling terms

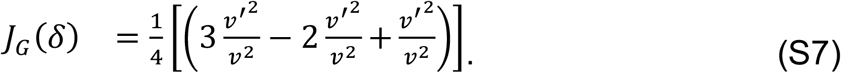

Simplifying yields a compact expression for the Fisher information in scalar mean-zero Gaussian-distributed random variables

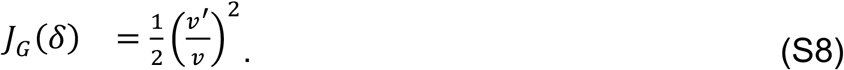

### Fisher information in mean-zero Laplace-distributed responses

The zero-mean Laplace distribution is given by

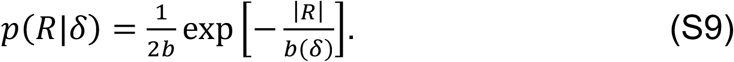

For receptive field responses *R* to be useful for discriminating a latent variable—here, fixation disparity—the conditional response distributions *p*(*R*|*δ*) must change with that latent variable. For a zero-mean Laplace distribution, this requirement entails that the scale parameter *b*(*δ*) is a function of the latent variable. The expression for the Fisher information that is contained in such Laplace-distributed responses is obtained by plugging the expression for the Laplace probability density (Eq. S1) into Eq. 1b in the main text, Specifically,

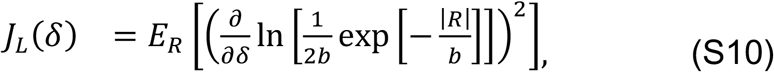

where, for notational simplicity, the dependence of the scale parameter on the latent variable has been dropped.

Using the product rule for logarithms and distributing the derivative operator

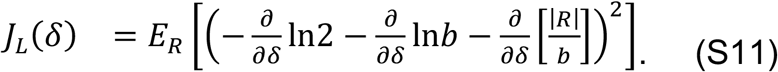

Taking the derivative

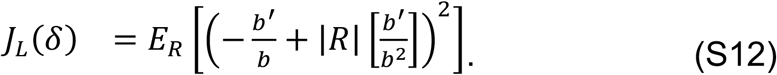

Expanding the square

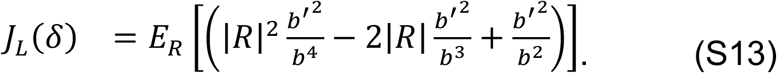

If *R* is mean-zero and Laplace-distributed, *E*[|*R*|^2^] *=* 2*b*^2^ and *E*[|*R*|] *= b*. Substituting and making use of the fact that *E*[*ax*] *= aE*[*x*] gives

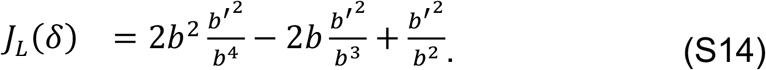

Simplifying

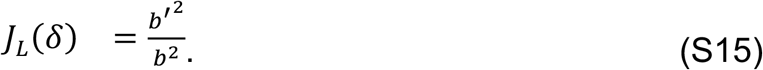

The standard deviation of the Laplace distribution is given by 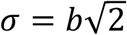, so the Fisher information can be expressed in terms of the response standard deviations and their derivatives with respect to disparity

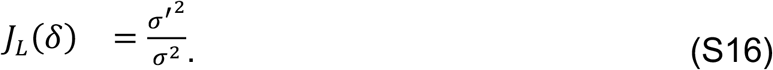

The Fisher information can also be expressed in terms of the variance. Rewriting,

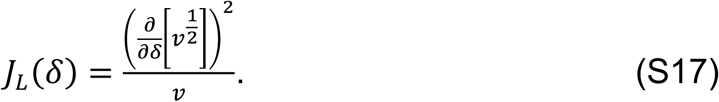

Taking the derivative

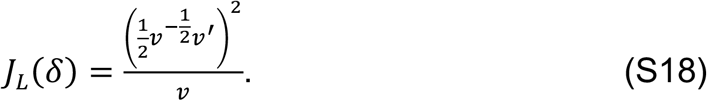

Simplifying yields an equivalent expression for the Fisher information in a scalar mean-zero Laplace-distributed random variable, expressed in terms of the variance

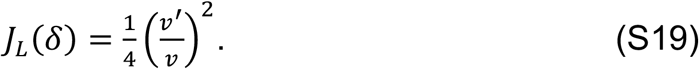

### Fisher information in multivariate mean-zero Gaussian-distributed Responses

The zero-mean Gaussian distribution is given by

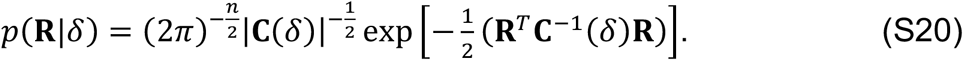

For the receptive field population response **R** to be useful for discriminating a latent variable—here, fixation disparity—the conditional response distributions *p*(**R**|*δ*) must change with that latent variable. For a zero-mean Gaussian distribution, this requirement entails that covariance **C**(*δ*) changes with the latent variable. By definition, the covariance matrices are positive definite and symmetric. The expression for the Fisher information that is contained in such Gaussian-distributed responses is obtained by plugging the expression for the Gaussian probability density (Eq. S20) into Eq. 1b in the main text.

Specifically,

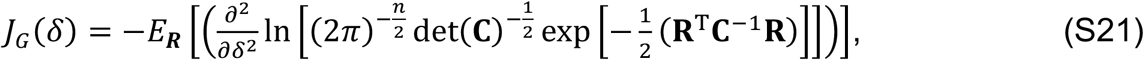

where, for notational simplicity, the dependence of the covariance matrix on the latent variable has been dropped.

Taking the log and distributing one of the two derivative operators

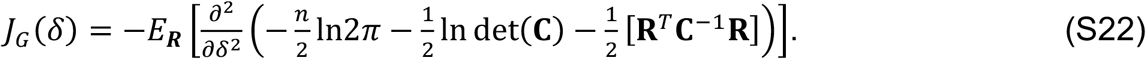

Taking the first derivative of each term

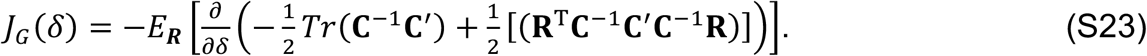

Taking the second derivatives, using the chain rule, the product rule, and the fact that the derivative of the trace is the trace of the derivative—that is, 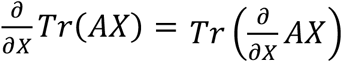—gives

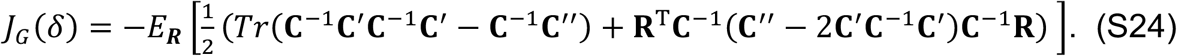

Rewriting using the cyclic property of traces and the fact that a scalar equals the trace of a scalar gives

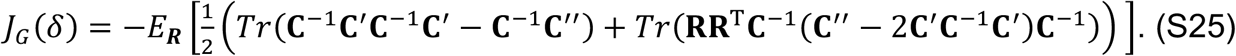

The expectation of the trace is the trace of the expectation and the expectation of the outer product of the mean-zero responses *E*_***R***_[**RR**^***T***^] equals the covariance matrix. It follows that *E*_***R***_[*Tr*(**RR**^***T***^**C**^-1(^**A**)] *= Tr*(*E*_***R***_[**RR**^***T***^]**C**^-1(^**A**) *= Tr*(**CC**^-1(^**A**) *= Tr*(**IA**) *= Tr*(**A**).

Hence,

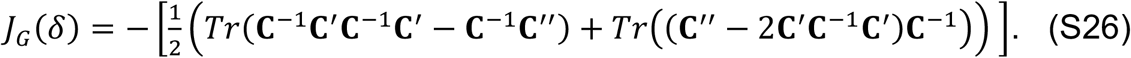

Distributing the trace operator gives

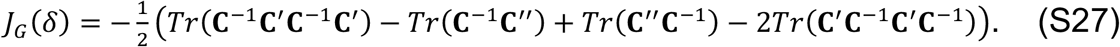

Subtracting like terms, and recognizing the cyclic property of traces (e.g. *Tr*(**ABC**) *= Tr*(**BCA**) *= Tr*(**CAB**)) gives

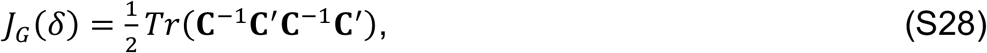

which is equal to Eq. 5 in the main text expression in the main text.

### Fisher information in multivariate mean-zero Laplace-distributed responses

For a zero-mean elliptical multivariate Laplace distribution the probability density is given by

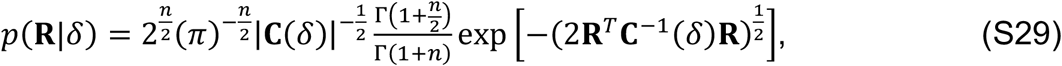

where *n* is the number of dimensions and Γ(.) is the gamma function, which interpolates the factorial function of the positive integers for the positive reals (Giller, 2005, 2024).

For a receptive field population response **R** to be useful for discriminating a latent variable—here, fixation disparity—the conditional response distributions *p*(**R**|*δ*) must change with that latent variable. For a mean-zero elliptically symmetric Laplace distribution, this requirement entails that covariance **C**(*δ*) changes with the latent variable. The expression for the Fisher information is obtained by substituting the expression for the Laplace distribution (Eq. S29) into Eq. 1 in the main text. Specifically,

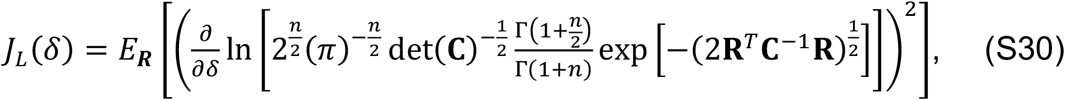

where, for notational simplicity, the dependence of the covariance matrix on the latent variable has been dropped.

Using the product rule for logarithms

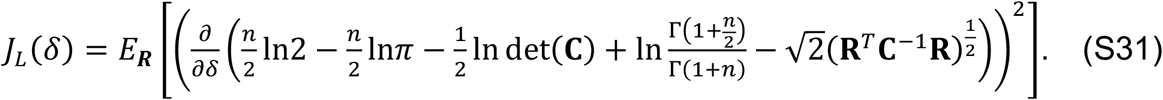

Taking the first derivative of each term and rearranging

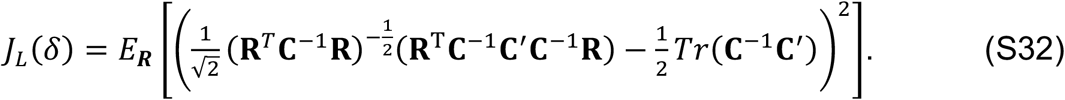

Taking the square

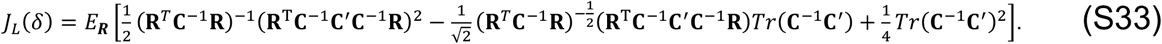

Rewriting for clarity

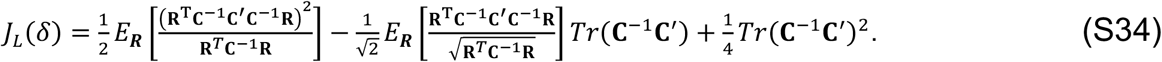

Substituting to reduce visual complexity

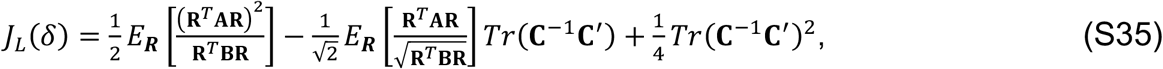

where **A** *=* **C**^-1^**C** ^′^**C**^-1(^ and **B** *=* **C**^-1^.

To evaluate the terms with expectation operators, we make use of results regarding the expectation of ratios of quadratic forms in elliptical random variables. Note that both terms in Eq. S35 with an expectation operator have the following form

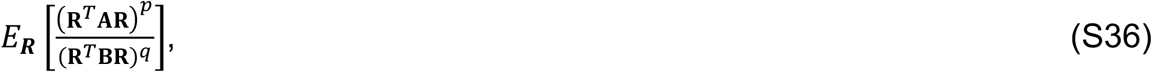

where *p =* 2 and *q =* 1 in the first term and *p =* 1 and 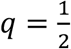 in the second term, so we seek a general expression for the expectation of this form. For arbitrary elliptical mean-zero random variable **R** ∼ E(**0**,**C**) we define 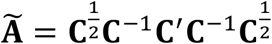 and 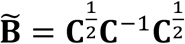 such that. 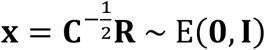 Substituting gives

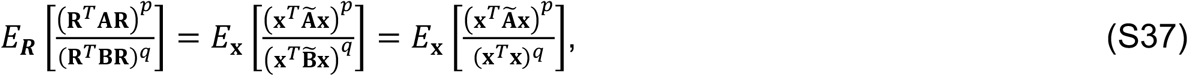

where the last equality holds because 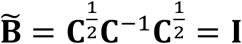.

The projection of a standard elliptical random variable (of arbitrary form) onto the unit sphere is independent of its vector magnitude (i.e. the magnitude of the projection equals 1.0). Expressing **x** *=* **u***w* where 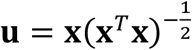 and 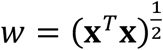 where **u** and *w* are independent of one another, allows one to rewrite Eq. S37 as

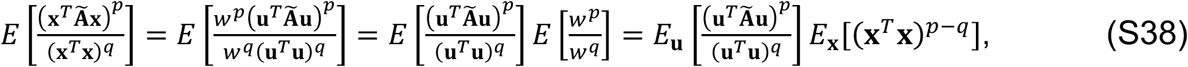

where the step separating the expectations follows from independence. Rewriting Eq. S38 using the fact that all random vectors projected onto the unit sphere are length 1.0 eliminates the denominator of the **u**-based term and gives

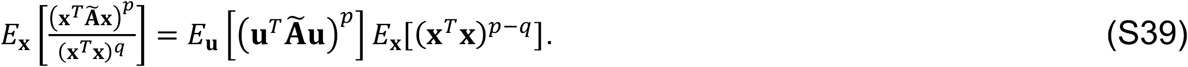

The solution of the p*r*oblem now reduces to finding expressions for each of the two expectations in Eq. S39 for the relevant values of *p* and *q*. The expression for the expectation *E*_**u**_[ (**u**^*T*^Ã**u**)^*p*^ ]is known for *p =* 2 and *p =* 1 when **u** is uniformly distributed on the unit sphere, as it is here. Specifically,

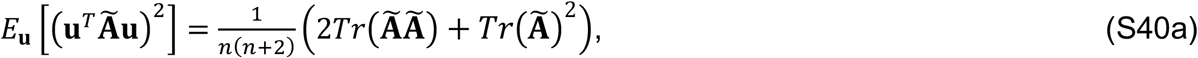

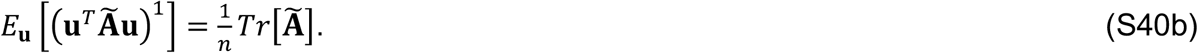

The expression for the expectation *E*_**x**_[(**x**^*T*^**x**)^*p*-*q*^] requires a bit more development. If the elliptical random variable **x** is multivariate Laplace, it can be expressed as 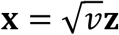, the product of independent standard scalar exponential random variable *v*∼Ex*p*(1) and standard multivariate normal random variable **z** ∼*N*(**0, I**).^1^ Therefore,

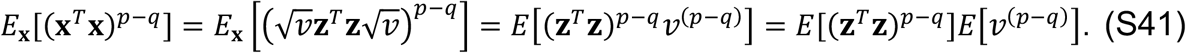

Both expectations in the term on the right-hand side are known. The first expectation on the right-hand side of Eq. S41 is the (*p* − *q*) moment of a chi-squared random variable with *n* degrees of freedom

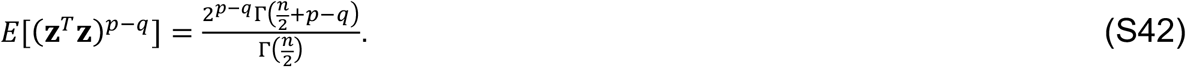

When *p =* 2 and *q =* 1, Eq. S42 simplifies to *n* by applying the identity Γ(*z* + 1) *= z*Γ(*z*).

When *p =* 1 and 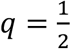, Eq. S42 equals 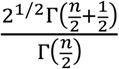

The second expectation on the right-hand side of Eq. S41 is the (*p* − *q*) moment of the exponential distribution, which is itself a scaled chi-squared 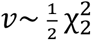 with two degrees of freedom

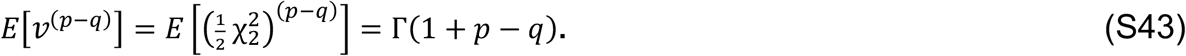

When *p =* 2 and *q =* 1, Eq. S43 evaluates to 1.

When *p =* 1 and 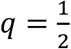, Eq. S43 evaluates to 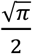.

Substituting Eqs. S42 and S43 into equation S41 gives

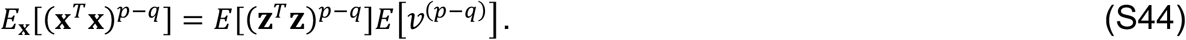

Substituting Eq. S44 into Eq. S39 gives

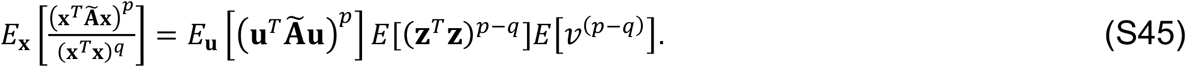

Evaluating Eq. S45 for *p* = 2 and *q* = 1, and for *p* = 1 and *q* = 0.5, by substituting in Eqs. S40ab, S42, and S43 gives

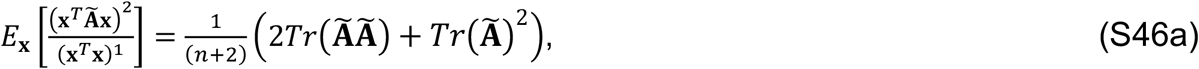

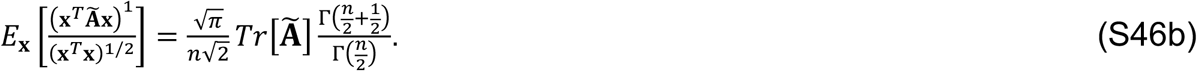

Rewriting the expression for Fisher information by substituting Eq. S37 into Eq. S35

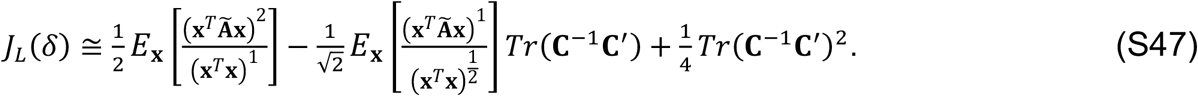

Substituting Eqs. S46ab into Eq. S47 gives

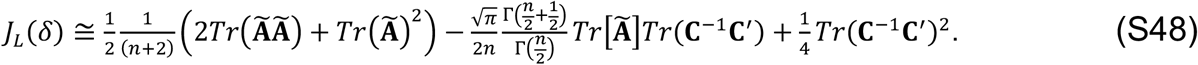

Re-expressing using the facts that 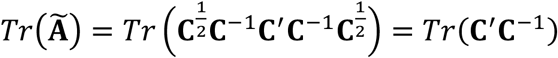 and that 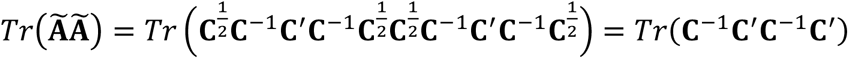.

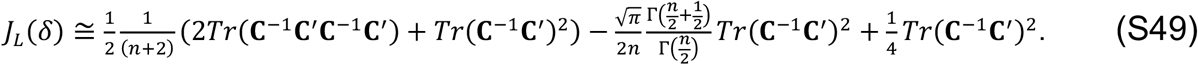

Grouping like terms

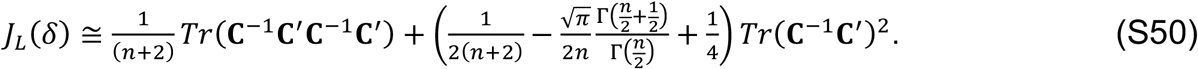

The second term tends to be small relative to the first term (especially for small *n*). Dropping the second term and rearranging the coefficient on the first term yields the approximation used in the main text

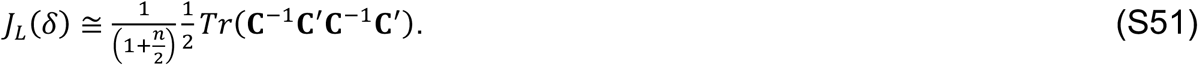

### The relationship between discrimination threshold and Fisher information

Here, we show that the relationship between Fisher information and the discrimination threshold is invariant to whether the estimator is unbiased or biased. The discrimination threshold is the critical stimulus separation

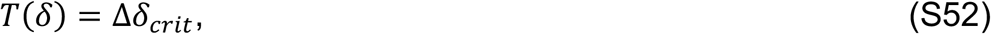

that corresponds to a criterion proportion correct in an *n*-interval (or *n*-look) 2AFC discrimination task. This criterion proportion correct is given by

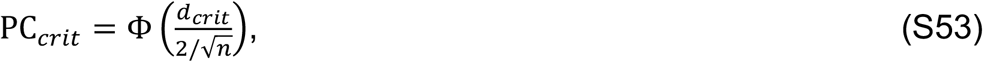

where Φ(.) is the standard cumulative normal, *d*_*crit*_ is the criterion sensitivity (or d-prime), and *n* is the number of intervals. For a criterion sensitivity of 1.0 in a two-interval task, for example, the criterion proportion correct is 0.76.

Sensitivity, or d-prime, for discriminating two different values of a latent variable is

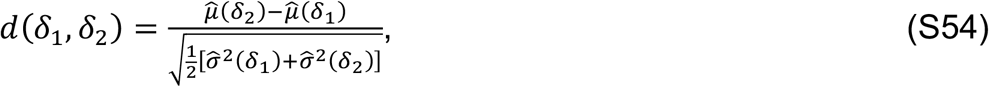

which is given by the distance between the mean estimates of the latent variable normalized by the square-root of the average variances of the estimates. The mean estimate 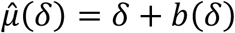 for an arbitrary value of the latent variable is given by its true value plus the bias.

For small differences △*δ* in the value of the latent variable, *d*-prime can be written as

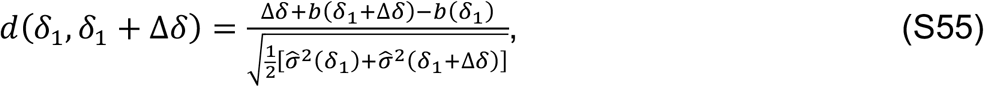

by substituting in expression for the mean estimate.

Assuming i) that 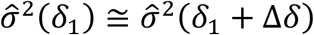, and ii) that the derivative of the bias is well approximated by 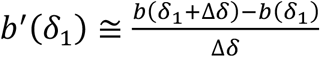 d-prime is approximately equal to

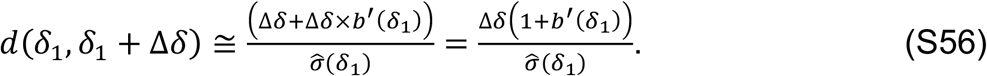

(These assumptions are not unreasonable for sufficiently small differences in the latent variable; recall that the definition of the derivative is 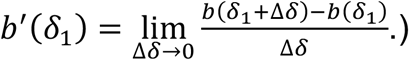

Substituting in the critical stimulus separation and criterion d-prime into Eq. S56 and multiplying through yields the following expression for the discrimination threshold

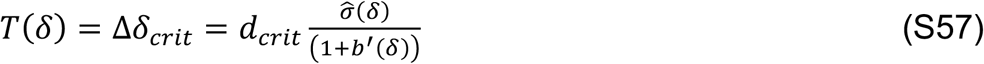

The Cramer-Rao bound on the variance of a biased estimator is given by

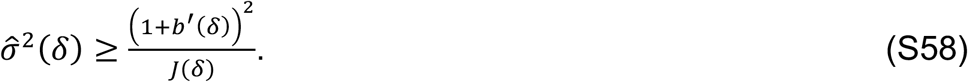

Dividing through Eq. S58 and taking the square-root yields

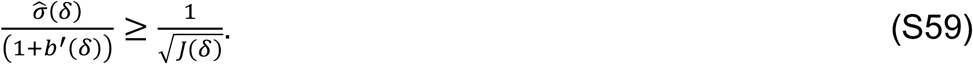

Substituting into Eq. S57 into Eq.S59 and rearranging yields the expression asserted in the main text

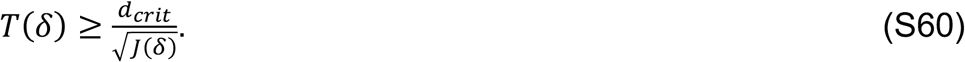

### Normalization via common binocular factor vs. individual monocular factors

Binocular receptive field responses could be normalized in at least two different stages of the binocular integration process. First, normalization could be carried out monocularly before binocular integration. In this scenario, the normalization factor is computed from the corresponding monocular contrast images. Second, the left- and right-eye contributions could be normalized jointly by a common factor, computed from the binocular contrast image. These two different approaches to normalization yield more different normalization factors when the left- and right-eye images are substantially different. Such differences occur more often with large fixation disparities and large disparity contrasts. But whether these potential differences in the normalization factors affect the response statistics relevant for disparity discrimination is unclear.

In the main text, normalization was carried out monocularly, before binocular integration (i.e. the first scenario). As a control, we redid the main analyses by normalizing the left- and right-eye response components by a common factor (i.e. the second scenario). The response statistics of binocular receptive fields remain largely unchanged at the granularity of the current analyses (see Fig. S1).

Specifically, in the control analysis, the broadband-normalized responses are given by

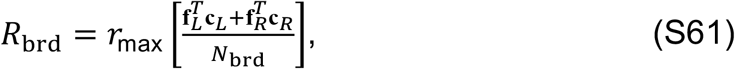

where the normalization factor is common to, or shared by, the two eyes. The common binocular broadband normalization factor is given by

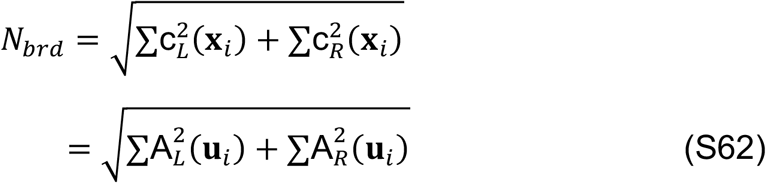

where 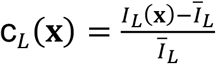 and 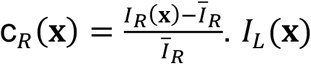 and *I*_*R*_(**x**) are intensity images for the left and right eye, respectively, and *Ī*_*L*_ and *Ī*_*R*_ are the corresponding average intensities for the left and right eye image, respectively.

Analogously, narrowband-normalized responses that are computed with a common binocular normalization factor are given by

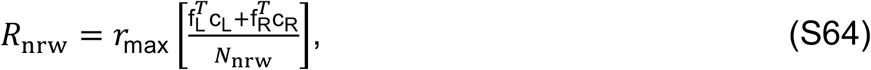

where common, binocular normalization factor is given by

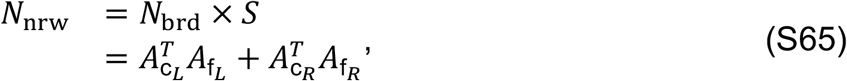

in which *S* is the cosine similarity between the binocular receptive field and the binocular image input (i.e., left and right eye images).

**Figure S1:**
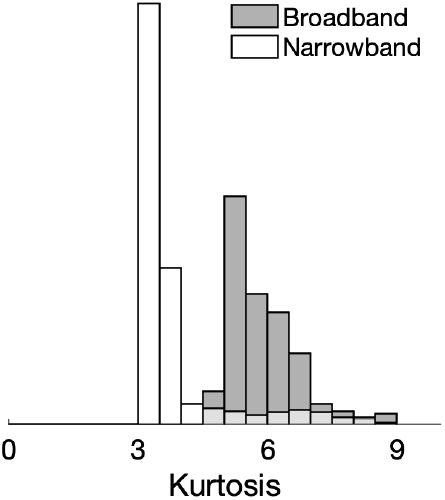
Histogram of kurtosis of normalized responses of all model binocular receptive fields with broadband (gray bars) and narrowband (white bars) normalization.

**Figure S2:**
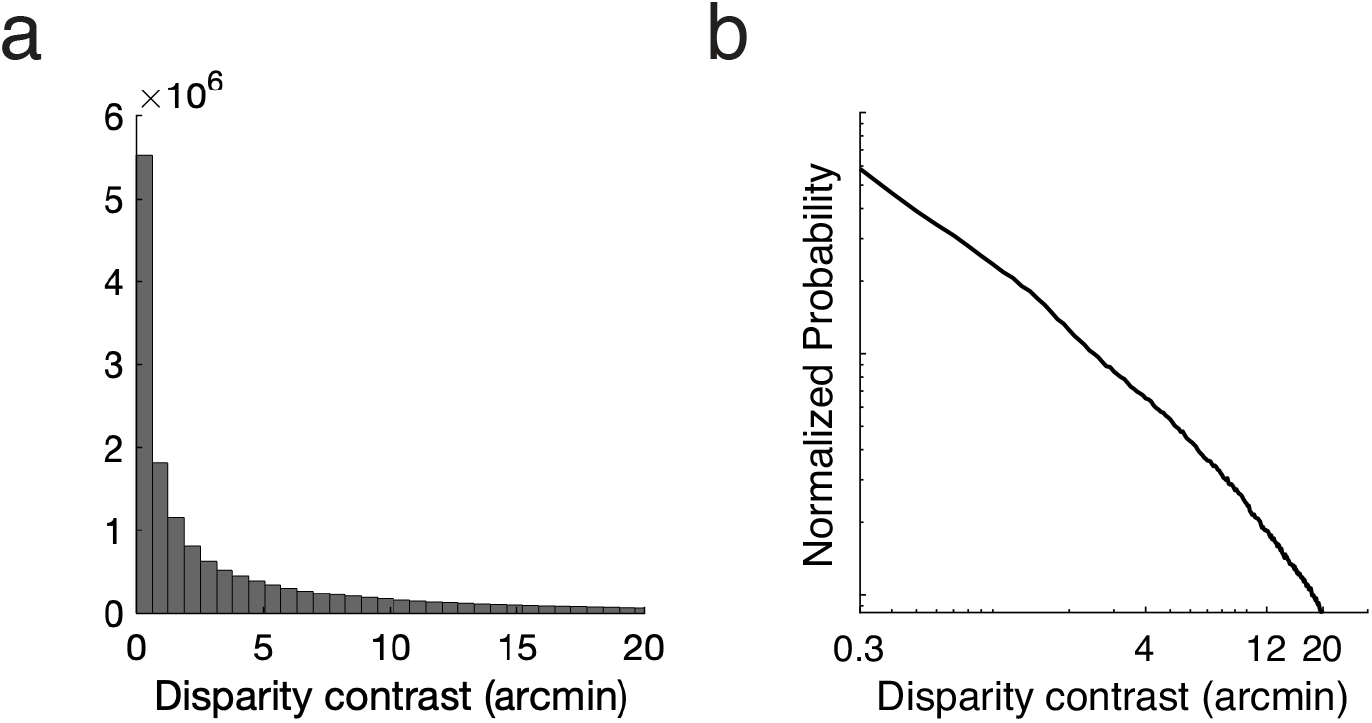
(a) The distribution of disparity contrasts computed from all stereo-image patches in the data set of size of 72 × 72 pixels (4/3ºx4/3º), sampled from 91 natural stereo-images. (b) Normalized probability of disparity contrast follows approximately a power law in the log domain.

## Footnotes

1 Note that there are several non-equivalent definitions of a multivariate Laplace probability density function (PDF). Eq. S29 defines a multivariate Laplace with Laplace-distributed radial components, whereas elliptical random variables that are constructed as an exponential scale mixture of a multivariate normal (see Eq. S41) yields a multivariate Laplace with Laplace-distributed marginals, and a less convenient analytic form for differentiation. Equations S47-S51 are therefore approximations.

## References

Adelson, E. H., & Bergen, J. R. (1985). Spatiotemporal energy models for the perception of motion. Journal of the Optical Society of America-A, 2(2), 284–299.

Anzai A., Ohzawa I., Freeman R.D. (1999). Neural mechanisms for encoding binocular disparity: receptive field position versus phase. Journal of Neurophysiology. 82:874–90.

Badcock, D. R., Schor, C. M. (1985). Depth increment detection function for individual spatial channels. Journal of the Optical Society of America-A, 2(7), 1211–1215.

Banks, M.S., Gepshtein, S., Michael S Landy, M.S. 2004. Why is spatial stereoresolution so low? Journal of Neuroscience 24 (9): 2077–89.

Blakemore, C. (1970). The range and scope of binocular depth discrimination in man. Journal of Physiology, 211(3), 599–622.

Burg, M.F., Cadena, S.A., Denfield, G.H. Walker, E.Y., Tolias, A.S., Bethge, M., Ecker, A.S. (2021). Learning Divisive Normalization in Primary Visual Cortex. PLOS Computational Biology, 17 (6): e1009028.

Burge, J. (2020). Image-Computable Ideal Observers for Tasks with Natural Stimuli. Annual Review of Vision Science, 6: 491–517.

Burge, J., and Geisler, W.S. (2011). Optimal Defocus Estimation in Individual Natural Images. Proceedings of the National Academy of Sciences, 108 (40): 16849–54.

Burge, J., Geisler, W.S. (2012). Optimal defocus estimates from individual images for autofocusing a digital camera. In: Proceedings of the IS&T/SPIE 47th annual meeting. Proceedings of SPIE. Burlingame, CA.

Burge, J., Geisler, W.S. (2014). Optimal disparity estimation in natural stereo images. Journal of Vision, 14(2):1, 1–18.

Burge, J., Geisler, W.S. (2015). Optimal speed estimation in natural image movies predicts human performance. Nature Communications, 6, 7900.

Burge, J., Jaini, P. (2017). Accuracy maximization analysis for sensory-perceptual tasks: Computational improvements, filter robustness, and coding advantages for scaled additive noise. PLoS Computational Biology, 13(2), e1005281.

Burge, J., McCann, B.C., and Geisler, W.S. (2016). Estimating 3D tilt from local image cues in natural scenes. Journal of Vision, 16 (13):2, 1–25.

Chin B.M., Burge, J. (2020). Predicting the partition of behavioral variability in speed perception with naturalistic stimuli. Journal of Neuroscience, 40:864–79.

Coen-Cagli, R., Dayan, P., Schwartz, O. (2012). Cortical surround interactions and perceptual salience via natural scene statistics. PLoS Computational Biology, 8:e1002405.

Coen-Cagli, R., Kohn, A., Schwartz, O. (2015). Flexible gating of contextual influences in natural vision. Nature Neuroscience, 18:1648–55.

Cook, R. D., Forzani, L. (2009). Likelihood-based sufficient dimension reduction. Journal of the American Statistical Association, 104(485), 197–208.

Cumming, B.G., DeAngelis, G.C. (2001). The physiology of stereopsis. Annual Review of Neuroscience, 24: 203–238.

Hecht, S., Shlaer, S., Pirenne, M.H. (1942). Energy, Quanta, and Vision. Journal of General Physiology, 25:819–840.

Heeger, D.J. Carandini, M. (2012). Normalization as a canonical neural computation. Nature Reviews Neuroscience, 13 (1): 51–62.

Carandini, M., Heeger, D.J., Movshon, J.A. (1997). Linearity and normalization in simple cells of the macaque primary visual cortex. Journal of Neuroscience, 17 (21): 8621–44.

De Valois, R.L, Albrecht, D.G., and Thorell, L.G. (1982). Spatial Frequency Selectivity of Cells in Macaque Visual Cortex. Vision Research, 22 (5): 545–59.

Fang, Z., Bloem, I.M., Olsson, C., Ma, W.J., Winawer, J. (2023) Normalization by orientation-tuned surround in human V1-V3. PLoS Computational Biology, 19(12): e1011704.

Fleet, D.J., Wagner, H., Heeger, D.J. (1996). Neural encoding of binocular disparity: energy models, position shifts and phase shifts. Vision Research, 36(12), 1839–1857.

Geisler, W.S. (1989). Sequential ideal-observer analysis of visual discriminations. Psychological Review, 96(2), 267–314.

Geisler, W.S. (2008). Visual perception and the statistical properties of natural scenes. Annual Review of Psychology, 59:167–92.

Geisler, W.S., and Ringach, D. (2009). Natural Systems Analysis. Visual Neuroscience, 26 (1): 1–3.

Giller, G. L. (2005). A generalized error distribution. Social Science Research Network, 2265027, 1–7.

Giller, G. L. (2024). An analytic solution for asset allocation with a multivariate Laplace distribution. Social Science Research Network, 4804682, 1–7.

Goris, R. L., Coen-Cagli, R., Miller, K. D., Priebe, N. J., & Lengyel, M. (2024). Response sub-additivity and variability quenching in visual cortex. Nature Reviews Neuroscience, 25(4), 237–252.

Haefner R.M., Bethge M. (2010). Evaluating neuronal codes for inference using Fisher information. Advances in Neural Information Processing Systems, vol. 23, pp. 1–9.

Herrera-Esposito D., Burge J. (2024). Optimal motion-in-depth estimation with natural stimuli. bioRxiv. 585059. 1–30.

Hou, C., Nicholas, S. C., Verghese, P. (2020). Contrast normalization accounts for binocular interactions in human striate and extra-striate visual cortex. Journal of Neuroscience, 40(13), 2753–2763.

Hubel, D. H., & Wiesel, T. N. (1962). Receptive fields, binocular interaction and functional architecture in the cat’s visual cortex. The Journal of Physiology, 160(1), 106–154.

Iyer A.V., Burge J. (2018). Depth variation and stereo processing tasks in natural scenes. Journal of Vision, 18(6):4. 1–22.

Iyer, A.V., Burge, J. (2019). The statistics of how natural images drive the responses of neurons. Journal of Vision, 19(13):4. 1–25.

Jaini, P., Burge, J. (2017). Linking normative models of natural tasks to descriptive models of neural response. Journal of Vision, 17(12):16. 1–26.

Mardia, K.V., Marshall, R.J. (1984). Maximum likelihood estimation of models for residual covariance in spatial regression. Biometrika, 71 (1): 135–46.

McFarland, J. M., Cui, Y., Butts, D. A. (2013). Inferring nonlinear neuronal computation based on physiologically plausible inputs. PLoS Computational Biology, 9(7), e1003143.

McKee, S.P., Levi, D.M., Bowne, S.F. (1990). The imprecision of stereopsis. Vision Research, 30(11), 1763–1779.

Mitchell, B.A., Carlson, B.M., Westerberg, J.A., Cox, M.A., Maier, A. (2023). A Role for Ocular Dominance in Binocular Integration. Current Biology, 33 (18): 3884–95.

Ni, L., & Burge, J. (2022). Encoding fidelity of binocular receptive fields with internal noise in the presence of external variability from natural scenes. Journal of Vision, 22(14), 4448–4448.

Olshausen, B.A., Field, D.J. (1996). Emergence of simple-cell receptive field properties by learning a sparse code for natural images. Nature, 381 (6583): 607–609.

Park, I. M., Archer, E. W., Priebe, N., & Pillow, J. W. (2013). Spectral methods for neural characterization using generalized quadratic models. Advances in Neural Information Processing Systems, vol. 26. 1–9.

Priebe, N.J., Mechler, F., Carandini, M., and Ferster, D. (2004). The Contribution of Spike Threshold to the Dichotomy of Cortical Simple and Complex Cells. Nature Neuroscience, 7 (10): 1113–1122.

Prince, S. J. D., Cumming, B. G., & Parker, A. (2002). Range and mechanism of encoding of horizontal disparity in macaque V1. Journal of Neurophysiology, 87(1), 209–221.

Qian, N., Zhu, Y. (1997). Physiological computation of binocular disparity. Vision Research, 37(13), 1811–1827.

Rieke, F., Warland, D., Van Steveninck, R.D.R., Bialek, W. (1997). Spikes: exploring the neural code. MIT press. Cambridge, MA.

Ringach, D.L. (2002). Spatial structure and symmetry of simple-cell receptive fields in macaque primary visual cortex. Journal of Neurophysiology 88 (1): 455–63.

Rieke, F., Warland, D., Van Steveninck, R. D. R., Bialek, W. (1997). Spikes: exploring the neural code. MIT press. Cambridge, MA.

Rust N.C., Schwartz O., Movshon J.A., Simoncelli, E.P. (2005). Spatiotemporal elements of macaque V1 receptive fields. Neuron, 46: 945–56.

Schor, C.M., Wood, I. (1983). Disparity range for local stereopsis as a function of luminance spatial frequency. Vision Research, 23(12), 1649–1654.

Schwartz, O., Simoncelli, E.P. (2001). Natural Signal Statistics and Sensory Gain Control. Nature Neuroscience, 4 (8): 819–825.

Sebastian S, Abrams J, Geisler WS. (2017). Constrained sampling experiments reveal principles of detection in natural scenes. Proceedings of the National Academy of Sciences, 114(28): E5731–E5740. doi:10.1073/pnas.1619487114.

Shadlen, M. N., Newsome, W. T. (2001). Neural basis of a perceptual decision in the parietal cortex (area LIP) of the rhesus monkey. Journal of neurophysiology, 86(4), 1916–1936.

Schwartz, O., Simoncelli, E.P. (2001). Natural Signal Statistics and Sensory Gain Control. Nature Neuroscience, 4 (8): 819–25.

Siderov, J., Harwerth, R.S. (1993). Precision of stereoscopic depth perception from double images. Vision Research, 33(11), 1553–1560.

Smallman, H.S., MacLeod, D.I. (1994). Size–disparity correlation in stereopsis at contrast threshold. Journal of the Optical Society of America-A, 11(8), 2169–2183

Stevenson, S. B., Cormack, L.K., Schor, C.M. (1989). Hyperacuity, superresolution, and gap resolution in human stereopsis. Vision Research, 29(11), 1597–1605.

Simoncelli, E.P., Olshausen, B.A. (2001). Natural Image Statistics and Neural Representation. Annual Review of Neuroscience, 24 (1): 1193–1216.

Skottun, B.C., De Valois, R.L., Grosof, D.H., Movshon, J.A., Albrecht, D.G., Bonds, A.B. (1991). Classifying Simple and Complex Cells on the Basis of Response Modulation. Vision Research, 31 (7-8): 1078–1086.

Tanabe S., Haefner R.M., Cumming, B.G. (2011). Suppressive mechanisms in monkey V1 help to solve the stereo correspondence problem. Journal of Neuroscience, 31:8295– 305.

Tyler C.W., Julesz, B. (1978). Binocular cross-correlation in time and space. Vision Research. 18:101–105.

Wainwright, M.J., Simoncelli, E.P. (2000). Scale Mixtures of Gaussians and the Statistics of Natural Images. In: Advances in Neural Information Processing Systems, vol. 12, pp. 855–61.

White, D.N., Burge, J. (2024). How distinct sources of nuisance variability in natural images and scenes limit human stereopsis. bioRxiv: 582383, 1–52. doi:10.1101/2024.02.27.582383.

## References

Giller, G. L. (2005). A generalized error distribution.

Giller, G. L. (2024). An Analytic Solution for Asset Allocation with a Multivariate Laplace Distribution. Available at SSRN 4804682.

